# Microalgal biofilm induces larval settlement in the model marine worm *Platynereis dumerilii*

**DOI:** 10.1101/2024.01.23.576855

**Authors:** Cameron Hird, Gáspár Jékely, Elizabeth A. Williams

## Abstract

A free-swimming larval stage features in many marine invertebrate life cycles. To transition to a seafloor-dwelling juvenile stage, larvae need to settle out of the plankton, guided by specific environmental cues that lead them to an ideal habitat for their future life on the seafloor. Although the marine annelid *Platynereis dumerilii* has been cultured in research labs since the 1950s and has a free-swimming larval stage, specific environmental cues that induce settlement in this nereid worm are yet to be identified. Here we demonstrate that microalgal biofilm is a key settlement cue for *P. dumerilii* larvae, inducing earlier onset of settlement, and enhancing subsequent juvenile growth as a primary food source. We tested the settlement response of *P. dumerilii* to 40 different strains of microalgae, predominantly diatom species, finding that *P. dumerilii* have species-specific preferences in their choice of settlement substrate. The most effective diatom species for inducing *P. dumerilii* larval settlement were benthic pennate species including *Grammatophora marina*, *Achnanthes brevipes*, and *Nitzschia ovalis*. The identification of specific environmental cues for *P. dumerilii* settlement enables a link between its ecology and the sensory and nervous system signalling that regulate larval behaviour and development. Incorporation of diatoms into *P. dumerilii* culture practices will improve the husbandry of this marine invertebrate model.

## 1. INTRODUCTION

Numerous marine invertebrates, from sponges to tunicates, face a crucial decision early in their life cycle - as a microscopic planktonic larva, they must select a suitable site on the sea floor on which to settle down, undergo metamorphosis from larval to juvenile form, and transition to a benthic, or bottom-dwelling, lifestyle. This major life cycle transition is guided by cues from the surrounding environment, which may attract or repel larvae from particular locations. Common cues that can induce settlement and metamorphosis in marine invertebrates include the presence of conspecifics, microbial biofilms, and potential food sources (1). An improved understanding of the process of larval settlement and metamorphosis is required, as this stage acts as a bottleneck in the life cycle and the process can shape adult marine invertebrate communities, in both aquaculture and biofouling scenarios, as well as in natural marine ecosystems predominated by invertebrates, such as coral reefs (2–4). Due to the diversity of invertebrate species and potential cues, as well as the challenge of studying microscopic interactions that occur in a macroscopic, three-dimensional environment, we still lack a comprehensive picture of the process of larval settlement, in terms of specific cue detection, the sensory modalities triggered within larvae, and the downstream internal signalling pathways that must be activated within a larva in order to drive metamorphosis, for any species.

The marine polychaete, *Platynereis dumerilii*, presents a model with good potential for gaining insight into the larval settlement process. *P. dumerilii* has been established as a laboratory model for several aspects of biology, including neurobiology, development, and evolution (5). *P. dumerilii* represents the relatively understudied phylum Lophotrochozoa,which also includes molluscs, bryozoans, brachiopods and platyhelminthes. Like many marine invertebrates, *P. dumerilii* has a life cycle that includes a planktonic larval stage that undergoes settlement and metamorphosis to a benthic juvenile. Understanding of sensory signalling processes in *P. dumerilli* larvae are greatly facilitated by molecular resources including extensive gene expression atlases in larval stages (6–11), a 3 day old full-body larval connectome (12), and knowledge of several nervous system signaling molecules and their receptors (13–15). In addition, genome editing tools have been successfully established in *P. dumerilii*, enabling the study of specific gene function through either gene knockout or knockdown (16,17).

Despite several decades of lab study, it remains unclear whether specific environmental settlement cues exist for *Platynereis dumerilii* larvae. In the natural environment, *P. dumerilii*, which are common in Mediterranean seas, but also distributed throughout European seas, are found in association with seagrass or macroalgae beds (5). However, *P. dumerilii* can be maintained in a laboratory environment throughout its life cycle, as long as juvenile and tube-dwelling adult stages are fed. The laboratory diet of *P. dumerilii* commonly consists of green microalgae, such as *Tetraselmis sp.* or *Spirulina sp.* either live, frozen, or dried (18–21). This is sometimes supplemented in later stages by a protein source such as *Artemia sp.* brine shrimp or rotifers, and shredded spinach leaves, which represent a macroalgae substitute. *P. dumerilii* are usually fed from approximately one week of age (19,20). This led us to question whether a potential food source may act as a natural settlement cue for *P. dumerilii* larvae? We hypothesized that if *P. dumerilii* would encounter such a cue earlier on in their development, even before they had developed the capacity to feed, they could transition out of their planktonic phase sooner, through a larval settlement process.

Previous studies of the diet of *Platynereis dumerilii* based on a natural population of worms from the Gulf of Naples (where lab cultures were originally sourced (5)) found that small worms primarily eat epiphytic microalgae growing on the macroalgal beds they inhabited, including diatoms and coccolithophores (22,23). Microbial marine biofilms are dominated by bacteria and diatoms (unicellular microalgae of the group Bacillariophyceae)(24). Microbial biofilms are a common inducer of larval settlement across many marine invertebrate phyla, including annelids (25,26)). Studies of larval settlement and metamorphosis in the polychaete *Hydroides elegans* previously identified both diatoms and bacteria as inductive cues for larval settlement (27–29). More recent studies have particularly emphasised a key role for bacteria as inducers of settlement and metamorphosis, through a variety of species-specific mechanisms, including a contractile injection system (30–32), and lipopolysaccharide-containing vesicles (33). These findings, together with the known dietary preferences of *P. dumerilii*, suggest that microbial biofilms may also play an inductive role in *P. dumerilii* larval settlement.

Here, we investigate microbial biofilms as potential larval settlement cues for *Platynereis dumerilii* larvae through a series of settlement assays. We base initial tests on biofilms developed from our adult culture boxes of *P. dumerilii*, then address individual components of biofilm, including bacteria, diatoms, green microalgae and coccolithophores, to determine from which source in the biofilm inductive cues may arise. Having established an important role for microalgae, particularly diatoms, in the induction of *P. dumerilii* settlement, we then ask whether this effect is species-specific. Finally, given the role of microalgae in larval settlement and its link to the early juvenile diet, we ask whether the inclusion of diatoms in the early diet of *P. dumerilii* lab cultures can enhance growth in their first month of culture.

## 2. MATERIALS AND METHODS

### Platynereis dumerilii culture

*Platynereis dumerilii* were cultured in the Marine Invertebrate Culture Unit (MICU) at the University of Exeter using a culturing method adapted from that described by Fischer and Dorresteijn (2004)(19). Adult cultures were maintained in the temperature-controlled (22 +/- 2 C) laboratories at the MICU, and cultured in 4 L styrene-acrylonitrile resin (SAN) boxes (Vitlab 36491), each containing 1.5 L of 1 micron-filtered and UV-sterilised artificial sea water (Tropic Marin Pro-Reef salt) at a salinity of 33 +/- 1 ppt, with a stocking density of 40 - 50 adult worms per box. Cultures were maintained under a 16:8 hr light:dark photoperiod, with a moon cycle simulated using 30 % light brightness overnight for 6 continuous nights within each 28-day cycle. Diet was varied with age, with larvae remaining unfed until 7 days post fertilization (dpf), at the point of box-culturing. Between 7 dpf and one month old, worms were fed with 10 mL mixed live microalgae, consisting primarily of *Tetraselmis suecica, Nannochloropsis salina* and *Phaeodactylum tricornutum*, pipetted into each box 3-times per week. Between 1 and 2 months old, worms were fed with 10 mL live rotifers per box 3-times per week, gut-loaded using RG Complete Microalgae (Reed Mariculture), separated from continuous cultures by sieving, and diluted in the same microalgal mix as the 7 dpf to one month old feed. Worms over 2 months old were fed using finely chopped (<3mm) frozen spinach, diluted with rotifers prepared as described above, at a concentration of 2.5 g of spinach in 5 mL rotifer preparation, with each box fed 5 mL of spinach solution 3-times per week. Water changes were performed every 2 weeks by 100 % replacement of the water with fresh artificial seawater. Excessive algal growth on the sides of the culture boxes was removed during the water change brushing with a dish brush.

Spawning was conducted by pipetting one male and one female epitoke using wide-aperture pasteur pipettes from culture boxes into a 100 mL glass beaker, containing approximately 80 mL fresh artificial seawater. Once spawning had taken place, adult worms were removed and the fertilised eggs left to settle for 10 minutes. After 10 minutes, the overlying water was poured off and the beaker refilled to approximately 80 mL with fresh artificial sea water, prior to being stored in an 18 °C incubator under a 16:8 hr light:dark light cycle for 7 days, with the excess jelly removed by pipetting and discarding approximately 10 mL from the bottom of each beaker at 24 hr post-fertilisation. At 7 dpf, larvae were cultured in 4 L SAN boxes, as outlined above, at an initial stocking density of 80-100 worms in 1.5 L artificial sea water.

Larvae used for the settlement assays were raised in 100ml glass beakers in an incubator on a 16:8 hr light/dark cycle at 18 °C as described above.

### Microalgae culture

Diatoms and other microalgae species were cultured in 50 ml vented flasks (BioLite 25cm^2^, ThermoFisher Scientific 130189) with 40 ml culture medium (autoclaved FSW containing F/2 nutrient medium at 1 ml per 2.5 L (ZM Systems), and silicate solution at 1 ml per 2.5L (ZM Systems) at 18°C in an incubator with fluorescent overhead illumination under a 16: 8 hr light:dark cycle.

Each strain was subcultured every 3 weeks by transferring 5-10 ml culture into a new flask and topping up to 40 ml with fresh culture medium. All work with microalgae cultures was carried out in a microbiological cabinet to prevent cross-contamination of cultures. For single-species microalgae biofilm assays, algae stock cultures were obtained through collaborators, purchase, or through an ASSEMBLE Plus Remote Access Grant from microalgal culture collections in the UK and Europe (Marine Biological Association (MBA) UK, Culture Collection of Algae and Protozoan (CCAP), Scotland, Roscoff Culture Collection (RCC), France, Stazione Zoologica Napoli (SZN), Italy, and the Belgian Coordinated Collection of Microorganisms (BCCM/DCG), Ghent. Further information on microalgal IDs and characteristics are provided in Supplementary data, Table S4. Note that microalgal cultures, although consisting predominantly of a single species, were not guaranteed to be axenic, and may have contained other microorganisms such as bacteria and ciliates, if these were present in original stock cultures provided.

### Preparation of biofilmed coverslips for initial settlement experiments

Six (3 for use in larval settlement assays, 3 for identifying bacterial community) 12 mm round glass coverslips (Roth P231.1) were placed in an adult *Platynereis dumerilii* culture box and left for 16 days to allow adequate time for a representative biofilm to grow. To retrieve the cover slips, they were collected in a sterile petri dish underwater and transported to the lab in 0.22 µm filtered seawater from the *P. dumerilii* culture box.

### Isolation of bacteria from *Platynereis* culture box biofilm

Methodology for isolation of bacteria from biofilm and settlement assays with single species of bacteria as biofilm was developed based on Freckelton *et al.* (2017)(33) and Lau *et al.* (2002)(34). In a laminar flow hood, biofilmed coverslips were rubbed with a sterile swab to transfer the biofilm into 10 ml autoclaved filtered seawater (FSW). For each coverslip, 200 µl of this 10 ml biofilm mix was plated undiluted, diluted 1:10 and diluted 1:100, with two replicate plates at each dilution, onto 1/2FSWt agar plates (filtered seawater tryptone ((35)), containing per litre 2.5 g tryptone, 1.5 g yeast extract, 1.5 ml glycerol, 15 g agar (1.5%) in filtered seawater, autoclaved). Plated biofilm cultures were allowed to grow overnight at 37°C, during which time multiple colonies developed on the plates. After 24 h, individual bacterial colonies were selected with a sterile autoclaved P10 pipette tip and transferred to a fresh 1/2FSWt agar plate. Isolated colony plates were left to grow at 37°C for half a day, then 4°C for 3 days. We selected colonies of interest based on differing gross morphology,with the aim of covering the morphological diversity of colonies growing on the plates, with 27 colonies selected overall. Single colonies were used to inoculate liquid cultures in 1/2FSWt broth, which were allowed to grow overnight. Despite repeated attempts, two of the isolated colonies grown on 1/2FSWt agar plates failed to generate liquid cultures and were therefore not used in subsequent settlement assays and sequencing ID. Liquid cultures were used to generate glycerol stocks for each of the 25 picked colonies; 0.5 ml 60% glycerol was combined with 0.5 ml bacterial culture and stock were stored at -80°C for subsequent experimentation.

### Identification of bacterial species isolated from *Platynereis* culture biofilm

Bacterial strains were streaked from -80°C glycerol stocks onto 1/2FSWt agar plates and grown overnight at 37°C. One loop of bacterial colony for each strain was suspended in 1 ml autoclaved MilliQ H2O, boiled for 15 minutes and centrifuged at 5000 x g for 2 minutes.

Supernatant was transferred to a new 1.5 ml eppendorf tube and saved as crude DNA extract for use as template in PCR reactions, stored at -20°C. PCR of the 16S ribosomal RNA gene was performed using bacterial universal primers ‘E.coli 8-26 26F’ (5’-AGAGTTTGATCCTGGCTCA-3’) and ‘E.coli 785-804 785R’ (5’-CTACCAGGGTATCTAATCC-3’). PCR conditions were: hold 95°C 5 min; 35 cycles 95°C 1 min, 55°C 1 min, 72°C 1 min; final extension 72°C 1 min. Each PCR reaction had a total volume of 20 µl, with final concentration 1X Phusion Buffer, 200 µM dNTPs, 0.5 µM primers, 0.02 U/µl Phusion polymerase (ThermoFisher Scientific F530L). PCR products were sequenced with sanger sequencing by Eurofins Genomics. Resulting sequences were BLASTed using BLASTn against microbial databases in NCBI to identify bacterial species (Supplementary data, Table S1).

### Preparation of coverslips with bacterial or microalgal biofilms for settlement assay

Initially round glass coverslips (Roth P231.1, 12 mm diameter) were used to prepare bacterial and microalgal biofilms for settlement assays, however due to the fragility of these, we transitioned to using round plastic coverslips. Plastic coverslips were prepared from A4 sheets of Aclar film (Agar Scientific AGL4458) with a 12.7 mm disc punch (Agar Scientific AGT5443). Discs were autoclaved to ensure sterility and subsequently coated with poly-D-lysine to improve adherence of the biofilm cells. For poly-D-lysine coating, coverslips were submerged in a solution of 10 µg/ml poly-D-lysine (Sigma-Aldrich P6407) at room temperature for 1 h, rinsed three times briefly in MilliQ H_2_O, then air-dried and stored at 4°C for up to 1 month before use in settlement assays.

Bacterial biofilms were generated from -80°C glycerol stocks streaked onto 1/2FSWt agar plates and incubated overnight at room temperature. Single colonies were used to inoculate 10 ml 1/2FSWt broth in 50 ml falcon tubes, which were incubated overnight at 37°C with shaking at 170 rpm. Cells were pelleted by centrifugation at 4°C for 30 min at 4000 x g and pellets were resuspended in 2 ml autoclaved FSW (0.22 µm filtered artificial seawater). Cell densities of resuspended pellets were calculated using OD_600_ values measured on a Nanodrop 8000 (ThermoFisher Scientific). Cell densities were adjusted to produce 10^8^ cells/ml for all strains. Cell suspensions were aliquoted into 24-well plates containing a round cover slip at 1.5 ml/well and incubated for 1 h at room temperature to allow bacterial surface attachment. Four replicates were performed for each bacterial strain and we tested five different bacterial strains, one representative strain from each of the five bacterial genera that we identified in the *Platynereis dumerilii* adult culture biofilm. After incubation, unattached bacteria were removed by gently dip-rinsing cover slips three times in 50 ml sterile artificial seawater.

To generate microalgae biofilms, cell suspensions of each microalgae strain were aliquoted into 24-well plates (ThermoFisher Scientific 930186 BioLite 6 well multidish) containing a round cover slip at 1.5 ml/well and incubated overnight at 18°C. Unattached microalgae cells were removed by gently dip-rinsing cover slips three times in 50 ml sterile seawater.

Negative control slips were also generated by incubating coverslips overnight with 1.5 ml diatom culture medium (artificial seawater with F/2 and silica) and dip-rinsing three times before transfer to 6-well plates for settlement assays. For tests of the effect of *Grammatophora marina* biofilm age, six coverslips were additionally incubated following dip-rinsing for 1 week, 2 weeks, and 1 month in the 24-well plates, with twice weekly change of culture medium to allow further development of the biofilm. We did not adjust microalgae culture cell density before addition to coverslips as cell density in the biofilm was expected to vary due to differences in the adherence and physical properties (e.g. size, shape) of different microalgae species. For each species, we calculated representative cell densities in the biofilm by imaging a 2.6 mm area for each of three coverslips per strain using a Zeiss Axiozoom v18 (Carl Zeiss GmbH) with a digital CMOS camera ORCA-Flash-4.0 (Hamamatsu) and 2.3X objective at 100X magnification. Example images of biofilms can be seen in Supplementary Material, Fig. S4, Fig. S8. Mean densities for each strain’s biofilm were calculated from these images using a custom Image J macro (available at https://github.com/MolMarSys-Lab/Platynereis-diatom_Hird_et_al_2024/macros) to calculate % area of cell coverage.

To test the inductive properties of the *Grammatophora marina* 1 day old biofilm, and the relative contribution of viable diatoms and other microbial cells, including bacteria, in the biofilm, we subjected biofilmed coverslips to the following treatments before use in settlement assays, with dip-rinsing before and after treatments: (1) 1 µm filtration of microalgae cell suspension (Millipore 1 µM filter SLFAO5010), to remove diatom cells but retain microbial cells, (2) submerged in 100% EtOH for 30 min, (3) submerged in 100°C seawater for 30 min, (4) submerged in 50°C seawater overnight, (5) dried at 50°C overnight, (6) treated with antibiotics (penicillin streptomycin, Sigma P4333, 5 units penicillin, 5 µg/ml streptomycin) at starting point of settlement assay, and (7) treated with antibiotics both prior to biofilm formation (10 units penicillin, 10 µg/ml streptomycin) and at starting point of settlement assay (5 units penicillin, 5 µg/ml streptomycin). To calculate viable cell densities in the treated biofilms, we imaged them on an Axiozoom v18 stereo microscope as described above, recording images under both white light and fluorescent light emission at 535 nm. Example images of biofilms can be seen in Supplementary Material, Fig. S4, Fig. S12.

### Larval settlement assays

Larval settlement assays were carried out in 6-well plates (ThermoFisher Scientific 130184 BioLite 6 well multidish) with 10 ml FSW and 30 *Platynereis dumerilii* nectochaete larvae per well. Larvae were sourced from a mixture of three different larval batches generated by crossing individual male and female *P. dumerilii* epitokes. Larvae were introduced to the 6-well plates containing biofilmed or negative control coverslips at the start of each experiment by pipetting with a P10 micropipette (Gilson). For the initial test of larval settlement in response to mixed biofilm developed from *P. dumerilii* adult boxes, three replicates of 30 three day old nectochaete larvae were tested and numbers of individuals crawling on coverslip were scored under a dissecting microscope every 12 h from 3.5 days to 5.5 days, and at 6.25 and 7.25 days after fertilization. To investigate larval settlement in response to different biofilm components, namely specific species of bacteria and microalgae, four replicates of 30 three day old nectochaete larvae were tested and numbers of individuals crawling on coverslips were scored at 3.5, 4, 5 and 6 dpf. To investigate more widely the species-specific settlement response of *P. dumerilii* to mono-species microalgal biofilms, with particular focus on diatom species, three replicates of 30 three day old nectochaete larvae were tested on three independent occasions, with 9 x 30 larvae tested overall per species. Numbers of larval settlement, death, and swimming larvae were scored as above at 24 and 48 h after the addition of larvae to the experimental plates.

For further investigations of the effects of mixed species biofilms and further characterization of the settlement response induced by the most effective diatom species, *Grammatophora marina*, additional settlement and growth assays were carried out. To test whether a mixed species biofilm could further increase settlement rate through synergistic effects, we carried out settlement assays comparing % larval settlement of 3 days old *Platynereis dumerilii* exposed to either single species biofilms of *G. marina* and *Achnanthes brevipes* or to various combinations of multiple species of microalgae. We tested four different microalgae mixtures: (1) a ‘settlement formula’ consisting of three species that induced high levels of settlement after 24 h as single species biofilms (*G. marina*, *A. brevipes*, *Nitzschia ovalis)*, (2) a ‘settlement formula’ consisting of six species that induced significant levels of settlement after 24 h as single species biofilms (*G. marina*, *A. brevipes*, *N. ovalis, Achnanthes sp., Chrysotila lamellosa, Fragilaria striatula)*, (3) a ‘growth formula’ consisting of six species that induced highest levels of growth after 11 days (*G. marina, N, ovalis, A. brevipes, Nitzschia laevis, Amphora sp.* (Roscoff Culture Collection RCC70623), *Achnanthes sp.*), and (4) a ‘growth formula’ consisting of 10 species that induced significant growth after 11 days (*G. marina, N, ovalis, A. brevipes, Nitzschia laevis, Amphora sp.* (Roscoff Culture Collection RCC70623), *Achnanthes sp.*, *C. lamellosa, F. striatula, Skeletonema dohrnii, Gomphonema sp.*). Biofilmed coverslips were prepared as above and allowed to develop overnight. For mixed biofilms, equal volumes of each individual species were pre-mixed in 15 ml falcon tubes before transfer to 24-well plates containing round cover slips. Numbers of larval settlement, death, and swimming larvae were scored as above at 24 and 48 h after the addition of larvae to the experimental 6-well plates. Six replicates of 30 larvae were tested and scored at 24 and 48 h. Alongside this, to determine the impact of biofilm age on larval settlement, we tested *P. dumerilii* larval settlement response to single species *G. marina* biofilms of different ages, with three replicates of 30 3 days old larvae scored at 24 and 48 h. Further investigation of the inductive properties of the one day-old *Grammatophora marina* biofilm was also carried out with six replicates of 30 larvae per control and treatment, and scoring of larval swimming, crawling, settlement on coverslip, and death, at 24 and 48 h after the addition of larvae. To investigate the influence of larval age on the settlement response induced by 1 day old *G. marina* biofilm, we also performed settlement assays with larvae introduced to a well with a biofilmed coverslip at 2, 3, 3.5, 4, 5, 6, 7 or 8 days old. Six replicates of 30 larvae were tested and scored at 24 and 48 h after induction for each age group. As previous *G. marina* treatment assays indicated a positive effect of antibiotic treatment on larval survival and settlement, we incorporated antibiotic treatment into these larval age assays, with penicillin-streptomycin (Sigma P4333, 5 units penicillin, 5 µg/ml streptomycin) added to the wells with larvae at the time of induction.

### Larval growth measurements

In addition to scoring larval settlement in assays at 24 and 48 h after induction, we measured the size of juveniles from settlement assays at 8 days after induction to investigate the effect of settlement biofilm on growth. For this, we maintained the larvae from settlement assays in their 6-well plates for 8 days after induction, exchanging 3 ml ASW every 2 days with a micropipette. We measured the body length and segment number of representative individuals by imaging individuals relaxed by drop-wise application of 1 M MgCl_2_ in ASW using a Zeiss Stemi 508 stereo microscope at 3X magnification with Zeiss Axiocam 208 colour camera and Zeiss Labscope software. Measurements of images were taken manually in Image J version 1.54f using the line tool and ‘Measure’ analysis. For each well of a 6-well settlement assay plate which initially contained 30 larvae, 6 representative 11 days old individuals were selected for measurement, based on their proximity to the centre of the coverslip in the well at the time of imaging. Size measurements were recorded for larvae from the testing of 40 different microalgae single species biofilms, and the testing of mixed versus single species biofilms. Example images of *Platynereis dumerilii* at 11 days old can be seen in Supplementary Material, Fig.S5, Fig. S9.

### Use of diatoms in large-scale culture of *Platynereis*

Given the positive effect of several microalgae species on the initial growth of *Platynereis dumerilii* larvae, we decided to test whether incorporating one or two diatom species into our regular culture of *P. dumerilii* could also enhance longer term growth on this larger scale. We focused on effects in the first 30 days of culture, as during this time period the worms are fed microalgae only, whereas from 30 days, the worms receive additional nutrition in the form of rotifers and spinach. Our aim in this case was not to test effects of the diatom species most effective for early settlement, but to establish whether addition of diatoms in general to the *P. dumerilii* diet could enhance growth longer-term. Diatom species chosen for this study were *Phaeodactylum tricornutum* (PLY100) and *Skeletonema dohrnii* (PLY612) due to their ability to be cultured easily on a large scale in our microalgal aquaculture setup, and their common and widespread use as food in several aquaculture environments (36). These were tested alongside common large-scale aquaculture strains of microalgae already established for marine invertebrate nutrition, including *Tetraselmis suecica* (CCAP66/4), *Nannochloropsis salina* (CCAP849/3), and *Isochrysis galbana* (CCAP927/20).

The large-scale culturing of marine microalgae at the MICU is performed in adapted autoclaved 1 L glass Duran bottles, with vented lids, containing a 6 mm silicon airline input, attached to a sterilised airstone. Low level aeration is provided constantly using a mains-powered air pump with 0.33 micron in-line air filter. Lighting is provided with red/blue LED light strips, mounted 15-30 cm from the culture vessels on an 18 hour light : 6 hour dark photoperiod. Microalgal cultures are established by addition of 2 mL axenic microalgae species stock to 500 mL culture medium (autoclaved FSW containing F/2 nutrient medium at 1 ml per 2.5 L (ZM Systems), and silicate solution at 1 ml per 2.5 L (ZM Systems)) for 7-14 days to establish prior to the first harvest. Once established, the aquaculture strains of microalgae are cultured in continuous culture by harvesting 50 % of the volume for feed 3 times per week, and refilling the vessel with fresh autoclaved culture medium. New continuous cultures are established once every 2 months.

Four microalgal feed treatments were established for testing: 1) a single non diatom species (*Tetraselmis suecica* or *Nannochloropsis salina*); 2) a single diatom species (*Phaedoactylum tricornutum* or *Skeletonema dohrnii*); 3) mixed species microalgal feed consisting of 3 species (*Tetraselmis suecica, Nannochloropsis salina* and *Phaeodactylum tricornutum*); 4) mixed species microalgal feed consisting of 5 species (*Tetraselmis suecica, Nannochloropsis salina, Isochrysis galbana, Phaedoactylum tricornutum* and *Skeletonema dohrnii*). For the single species treatments, worms were fed on one or the other of the two listed species, depending on which was in culture at the point of establishing that replicate. Mixed species treatments consisted of equal volumes of each component species. Worms for the culturing trials were maintained unfed in an incubator at 18°C until they reached 7 dpf, at which point they were cultured in 4 L SAN boxes containing 1.5 L ASW at a stocking density of 80 worms per box, established by individually pipetting worms over. Each box was assigned to one of the 4 aforementioned feeding regimes and fed by addition of 10 mL of the relevant microalgal mix, 3 times per week, commencing from 7 dpf. At 30 days old, a maximum of 10 worms per culture box were removed, immobilised by transferring them into a 1 M MgCl_2_ in ASW solution, and the number of segments of each worm counted under a Zeiss Stemi 508 stereo microscope and the 0-30 day growth rate established by dividing the total number of segments by 30 days.

### Statistical analyses

Data analysis and plotting was carried out in Python 3, using pandas and seaborn packages(37,38). Scripts and raw data files for data analysis and plots are available on Github at https://github.com/MolMarSys-Lab/Platynereis-diatom_Hird_et_al_2024.

Each data set was initially checked for normality using a Shapiro-Wilk test, which confirmed non-normal distributions. A Kruskal-Wallis test was used to assess significant differences in treatment and control in initial mixed biofilm experiments. For subsequent species-specific settlement assays, a Mann-Whitney U rank test was used to test for significant differences in both % larval settlement between each species and the negative control. A Bonferroni correction was applied to the p-value cutoff when comparing larval settlement across 40 different microalgae species (Fig. 2), to account for potential false positives with this higher sample number. Correlation between % larval settlement and biofilm % coverage was tested using Kendall’s tau test, both for all 40 microalgae species tested, and for the subset of microalgae found to be significantly inductive at at least one time point according to the Mann-Whitney U rank test with Bonferroni correction. We also applied Mann-Whitney U rank test with Bonferroni correction to investigate whether the % larvae dead at 24 and 48 h was significantly different in any microalgae species compared to the negative control.

A Kruskal-Wallis test with Dunn’s posthoc test was used to investigate whether the initial settlement responses of *Platynereis dumerilii* at 24 h varied according to specific features of microalgae including group, cell shape, cell assemblage and cell size. To assess whether encounter with the 40 species of algae led to significantly different growth by 11 days of age compared to the no biofilm control samples, we applied a Mann-Whitney U rank test with Bonferroni correction. Subsequent experiments were also analyzed for significant differences to the control using Mann-Whitney U rank test, and Kruskal Wallis test with Dunn’s posthoc test was used to check for significant differences between all potential pairwise comparisons. For each experiment, we recorded biofilm densities as described above and used these to check for any correlation with larval settlement rate using Kendall’s tau test.

## 3. RESULTS

### Mixed biofilm and components

Initial testing of the settlement rate of *Platynereis dumerilii* larvae in response to mixed microbial biofilms grown in adult worm culture boxes indicated that larvae settle earlier in the presence of a mixed microbial biofilm than they would in the absence of this environmental cue (Fig. 1B). Numbers of settled larvae peaked at a maximum of ∼80% larval settlement on coverslips with biofilm at 36 h after induction. On control coverslips lacking a biofilm, a significantly lower maximum of 30% larval settlement occurred at 48 h into the assay.

**Figure 1.**
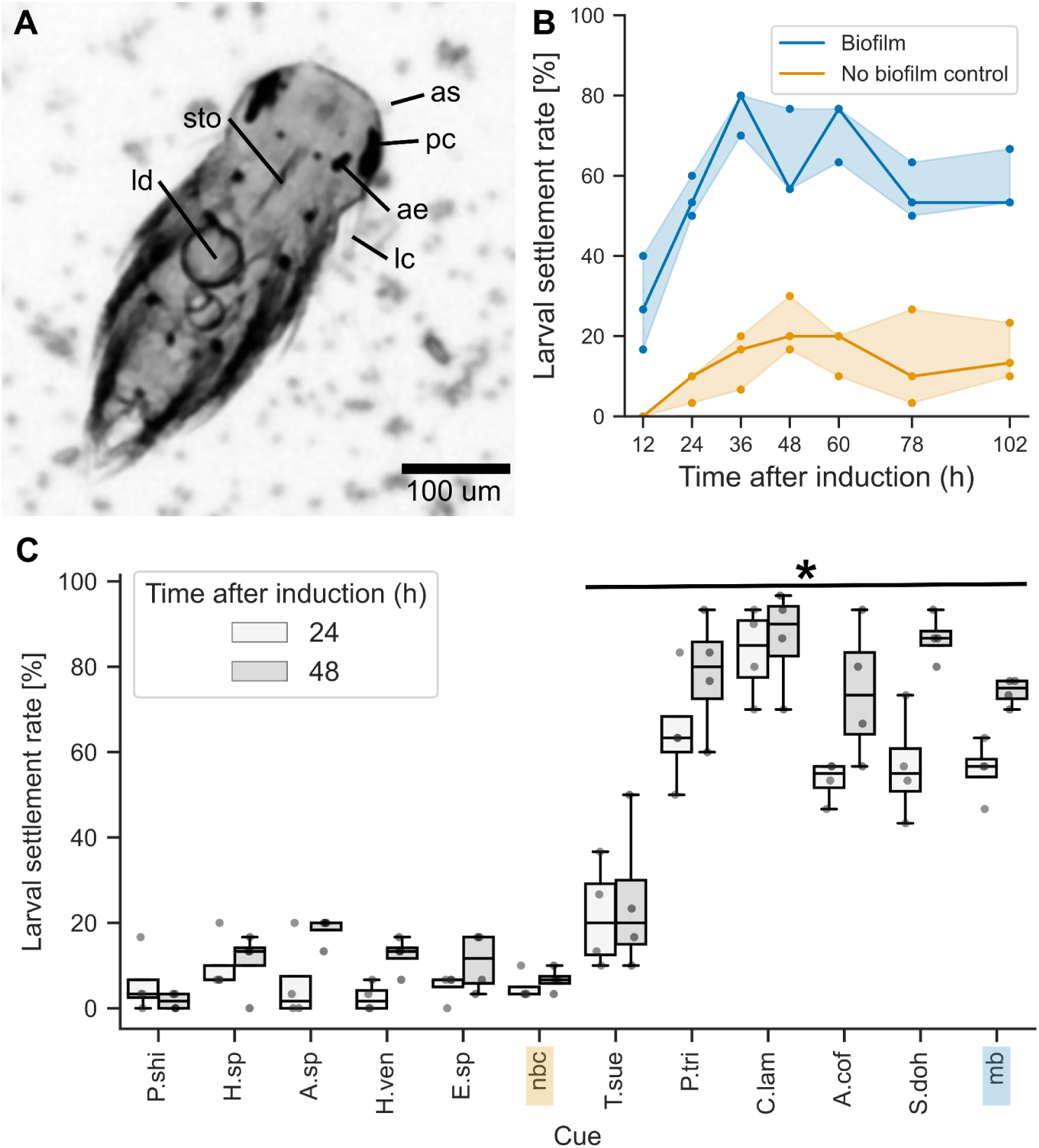
Microalgae, not bacteria, are the primary inducer of *Platynereis dumerilii* larval settlement in a mixed microbial biofilm. (A) Light micrograph of 3.5 days old *P. dumerilii* larva settled on a biofilm containing microalgae (ld = lipid droplet, sto = stomodeum, as = antennal stub, pc = pigment cell, ae = adult eye, lc = lateral cirrus). (B) Line plot of median % larval settlement over time. Line shadows indicate 95% confidence interval. Dot points indicate raw data. Kruskal-Wallis Test finds significantly higher % settlement in the presence of a mixed biofilm (p-value = 1.04E-07). (C) Grouped box plot of % larval settlement in response to different bacterial or microalgal biofilms after 24 h and 48 h exposure. No biofilm negative control and mixed biofilm positive control are highlighted in orange and blue respectively. * = p<0.05 at both 24 and 48 h after induction (Mann-Whitney U test). P.shi = *Pseudoalteromonas shioyasakiensis*, H.sp = *Halobacillus sp. G-12*, A.sp = *Alteromonas sp. strain JLT1934*, H.ven = *Halomonas venusta*, E.sp = *Exiguobacterium sp. WR-24*, nbc = no biofilm control, T.sue = *Tetraselmis suecica*, P.tri = *Phaeodactylum tricornutum*, C.lam = *Chrysotila lamellosa*, A.cof = *Amphora coffeaeformis*, S.doh = *Skeletonema dohrnii*, mb = mixed biofilm.

Significantly higher numbers of larvae settled on biofilmed coverslips were maintained throughout the settlement assay, up to 102 h (> 4 days) after initial induction at 3 days of age. Subsequent testing of the effects of individual bacterial species isolated from the biofilm, and individual species of microalgae including green microalgae (*Tetraselmis suecica*), coccolithophores (*Chrysotila lamellosa*) and diatoms (*Phaeodactylum tricornutum*, *Skeletonema dohrnii* and *Amphora coffeaeformis*) showed that bacterial biofilms alone did not induce significant levels of *Platynereis* settlement, but biofilms of individual microalgae species did induce significantly higher rates of settlement compared to the control coverslip which lacked a biofilm (Fig. 1C). Unlike the green microalgae *T. suecica*, which induced relatively low levels of settlement, single species diatom and coccolithophore biofilms were capable of inducing equal or greater numbers of larval settlement as the original mixed biofilm sourced from the adult worm culture boxes.

### Single-species microalgal biofilm

Having established that the primary inductive component in a mixed microbial biofilm was likely of microalgal origin, we then proceeded to test a number of additional different single species biofilms in settlement assays, with additional replication across multiple batches of *Platynereis dumerilii* larvae, to determine whether the settlement response of *P. dumerilii* to microalgae was species-specific. Of 40 species tested, we found that the species which induced the greatest median settlement after both 24 and 48 h was the chain-forming diatom *Grammatophora marina*. According to a Mann-Whitney U rank test with Bonferroni correction, of the 40 microalgae species tested, 13 species induced significantly greater larval settlement after 24 h compared to a no biofilm control, with 10 of these 13 species maintaining significantly greater levels of larval settlement 48 h into the assay (Fig. 2A, B). Apart from *G. marina*, other highly inductive microalgae species were *Nitzschia* species, including *Nitzschia ovalis*, *N. laevis* and *N. epithemoides*, and *Achnanthes* species, including *Achnanthes sp.*, and *A. brevipes*. Larval settlement response was also species-specific in that different microalgae species from the same genus elicited significantly different levels of *P. dumerilii* larval settlement. For example, two strains of *Achnanthes yaquinensis* tested did not induce significant larval settlement at 24 and 48h, while *Achnanthes sp.* and *A. brevipes* did.

**Figure 2.**
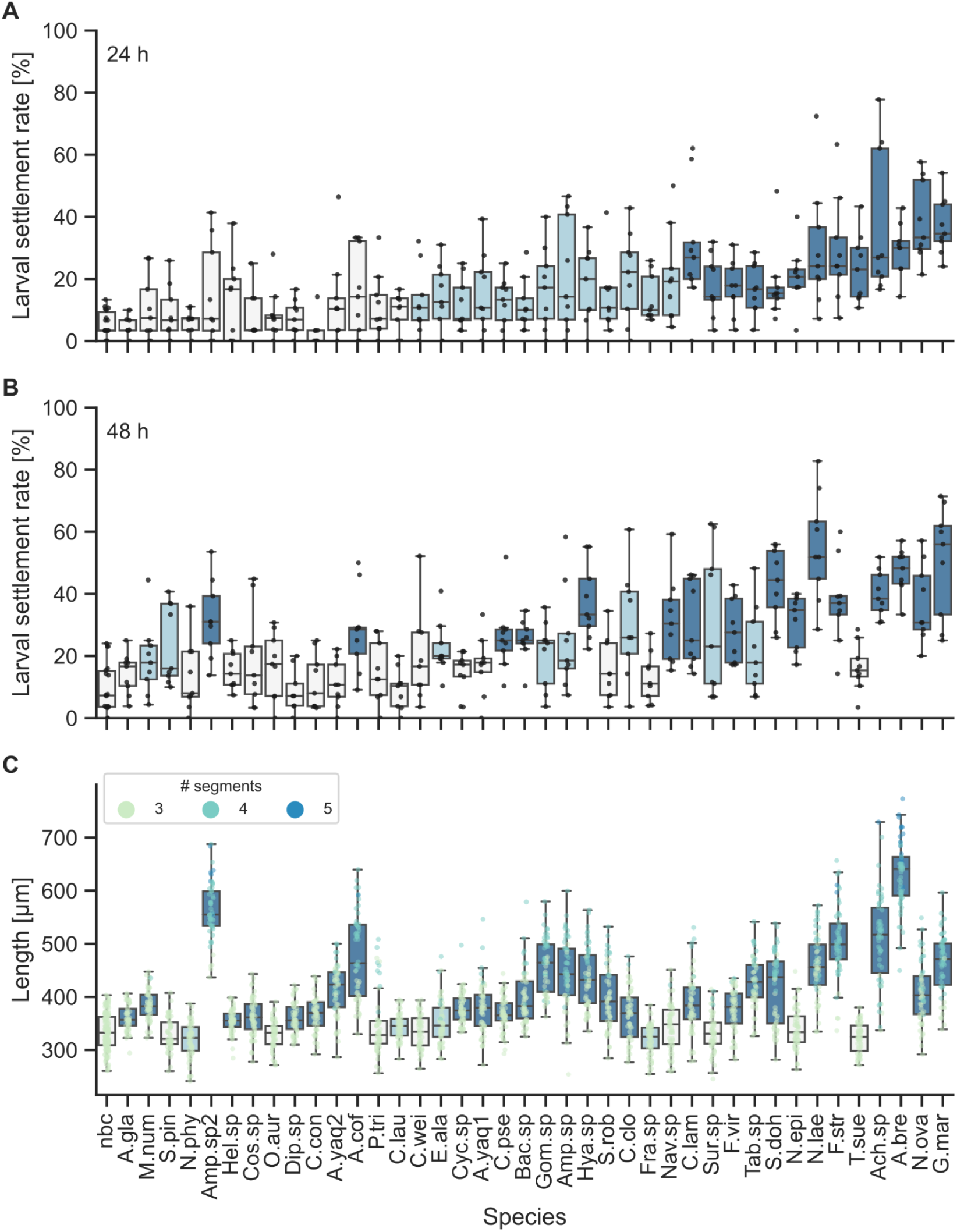
*Platynereis dumerilii* show a species-specific response to microalgae biofilms as inductive cues for settlement. Box plots with scatter plot overlay of (A) % larval settlement in response to different monospecies microalgal biofilms after 24 h and (B) 48 h exposure, and (C) total length of individual larvae exposed to the same biofilms for 8 days. Scatterplot points in (C) are coloured by segment number of each larva. Boxplots are coloured according to the p-value result of a Mann-Whitney U rank test of % larval settlement at 24 and 48 h, or total length at 8 days, in each species versus the no biofilm control; p>0.05 = pale grey, p<0.05>0.00125 = light blue, p>0.00125 = dark blue. A.gla = *Asterionellopsis cf glacialis*, M.num = *Melosira nummuloides*, S.pin = *Staurosirella pinnata*, N.phy = *Navicula phyllepta*, Amp.sp2 = *Amphora sp RCC7063*, Hel.sp = *Helicotheca sp.*, Cos.sp = *Coscinodiscus sp.*, O.aur = *Odontella aurita*, Dip.sp = *Diploneis sp.*, C.con = *Chaetoceros convolutus,* A.yaq2 = *Achnanthes yaquinensis DCG0053*, A.cof = *Amphora coffeaeformis*, P.tri = *Phaeodactylum tricornutum*, C.lau = *Chaetoceros cf lauderi*, C.wei = *Conticribra weisflogii*, E.ala = *Entomoneis alata*, Cyc.sp = *Cyclotella sp.*, A.yaq1 = *Achnanthes yaquinensis DCG0020*, C.pse = *Chaetoceros pseudocurvisetus*, Bac.sp = *Bacteriastrum sp.*, Gom.sp = *Gomphonema sp.,* Amp.sp = *Amphora sp.*, Hya.sp = *Hyalosira sp.*, S.rob = *Seminavis robusta*, C.clo = *Cylindrotheca closterium*, Fra.sp = *Fragilariopsis sp.*, Nav.sp = *Navicula sp.*, C.lam = *Chrysotila lamellosa*, Sur.sp = *Surirella sp.*, F.vir = *Fragilariformia virescens*, Tab.sp = *Tabularia sp.*, S.doh = *Skeletonema dohrnii*, N.epi = *Nitzschia epithemoides*, N.lae = *Nitzschia laevis*, F.str = *Fragilaria striatula*, T.sue = *Tetraselmis suecica*, Ach.sp = *Achnanthes sp.*, A.bre = *Achnanthes brevipes*, N.ova = *Nitzschia ovalis*, G.mar = *Grammatophora marina*, nbc = no biofilm control.

Based on the results of Kruskal-Wallis tests with Dunn’s posthoc testing, the initial settlement responses of *Platynereis* larvae to different microalgal species varied significantly depending on the cell shape, size, and phylogenetic group of the species tested (Supplementary data, Fig. S3). Round or centric diatom species induced significantly lower levels of larval settlement overall. The results of these tests indicate that the settlement response of *Platynereis* larvae is most strongly induced by pennate chain-forming diatoms under 50 micron in length. According to a Mann-Whitney U rank test with Bonferroni correction, no microalgae strains caused significantly more larval death than the control scenario (Supplementary data, Fig.S1, Table S6). Comparing % larval settlement with the % biofilm coverage of each microalgae species tested using Kendall’s rank correlation, we found a significant but negligible positive correlation between biofilm coverage and the rate of larval settlement (Supplementary Data, Fig.S2), leading us to conclude that biofilm coverage is not of primary importance to predicting settlement rate.

Although *Grammatophora marina* induced the highest median settlement rates in *Platynereis dumerilii* larvae at 24 and 48 h after induction, biofilms of this species did not support the highest median growth over the week following settlement. Of the 40 strains tested, 28 induced significantly more growth by 11 days compared to a no biofilm control (Fig. 2C). Top inducers of initial settlement, *G. marina*, *N. ovalis* and *A. brevipes* all induced significant growth after 11 days compared to control. Although not an inducer of significantly higher early settlement, *Amphora sp RCC7063* did induce significant growth by 11 days, and was one of the few species in which we found juveniles with 5 segments present. Other species that enabled growth of juvenile *Platynereis* to 5 segments by 11 days were *Amphora coffeaeformis*, *Fragilaria striatula*, *Achnanthes brevipes* and *Achnanthes sp*.

### Biofilm age and composition

In subsequent settlement and growth assays, we investigated the effects of biofilm age and composition on larval settlement rates in *Platynereis dumerilii*. Here, we focused primarily on *Grammatophora marina* biofilms, as this species induced the highest median % larval settlement in previous assays. Overall, the age of *G. marina* biofilm did not significantly alter the settlement rates of *P. dumerilii* larvae over 48 h, or the growth achieved by 11 days (Fig. 3, Supplementary data, Fig. S7., Table S9, S10). Similarly, although mixed species biofilms induced significantly greater larval settlement at 24 h compared to the no biofilm control, they did not cause significantly greater larval settlement at 24 or 48 h compared to single species biofilms of either *G. marina* or *Achnanthes brevipes.* However, mixtures of 3 diatom species selected based on their positive effects on larval settlement in initial testing, or 6 diatom species selected for their positive effects on larval growth in initial testing, did induce significantly more growth in larvae by 11 days compared to larvae exposed to either a single species *G. marina* biofilm, a mixed biofilm of 6 species selected for positive effects on settlement, or a mixed biofilm of 10 species selected for their positive effects on settlement and growth. Growth by 11 days in larvae exposed to the single species *A. brevipes* biofilm was equivalent to that shown by larvae in the best mixed species biofilms. As with the initial single species testing, biofilms tested in the biofilm age and composition settlement assays did not cause significantly greater rates of larval death (Supplementary data, Fig.S6, Table S9), and there was no significant correlation between the density of biofilms tested and larval settlement rates (Supplementary data, Fig. S7).

**Figure 3.**
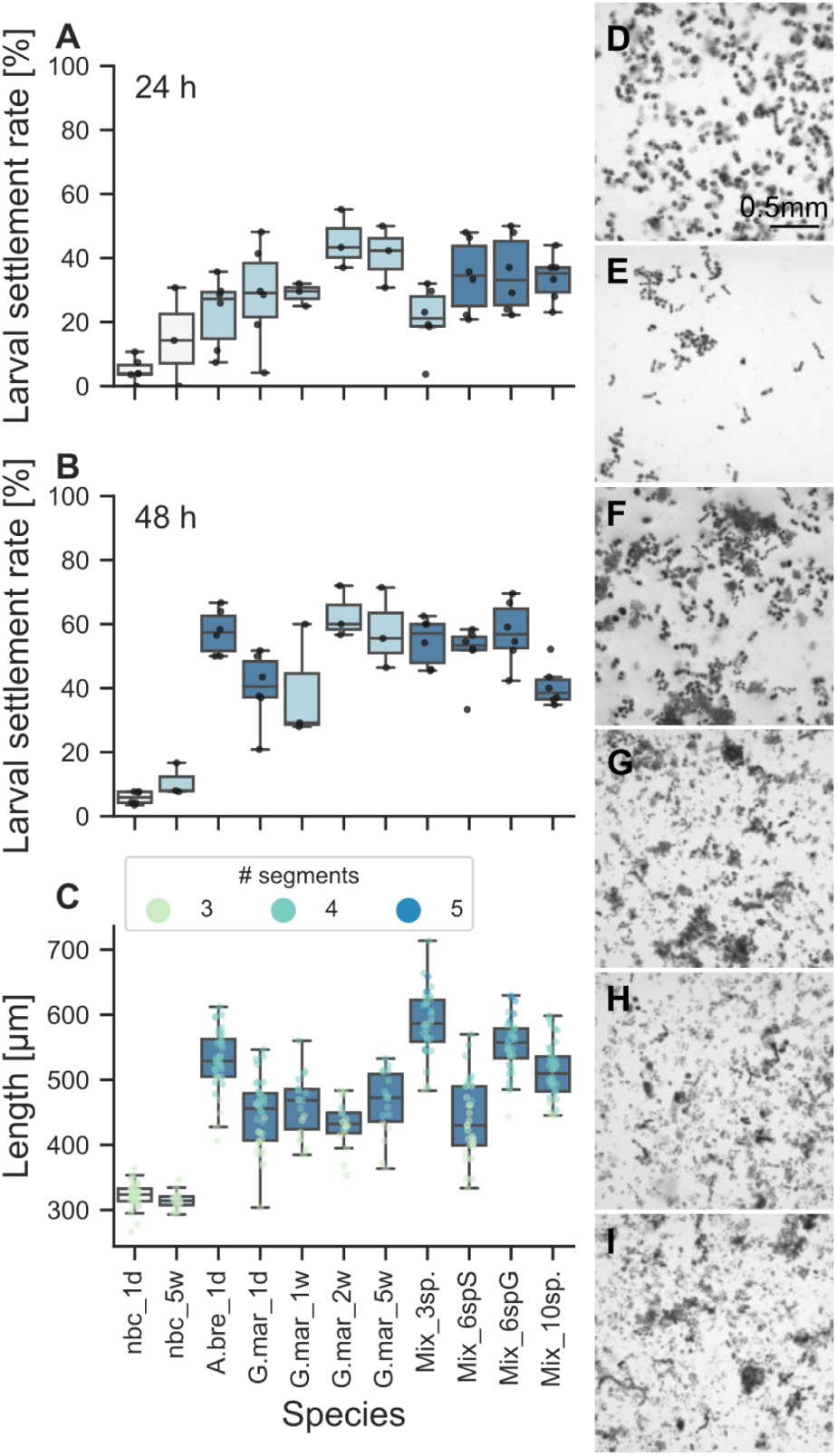
Impact of biofilm age and composition on *Platynereis dumerilii* larval settlement and early growth. Box plots with scatter plot overlay of (A) % larval settlement in response to different monospecies or mixed microalgal biofilms after 24 h and (B) 48 h exposure, and (C) total length of individual larvae exposed to the same biofilms for 8 days. Scatterplot points in (C) are coloured by segment number of each larva. Boxplots are coloured according to the p-value result of a Mann-Whitney U rank test of % larval settlement at 24 and 48h, or total length at 8 days, in each species versus the no biofilm control; p>0.05 = pale grey, p<0.05>0.005 = light blue, p>0.005 = dark blue. (D - I) Light micrograph example images of different monospecies and mixed biofilms used in settlement assays. (D) *Achnanthes brevipes*, (E) *Grammatophora marina*, (F) 3 species mixed biofilm, (G) 6 species mixed biofilm, targeted for induction of early settlement, (H) 6 species mixed biofilm, targeted for induction of both early settlement and longer term growth, (I) 10 species mixed biofilm. A.bre_1d = *A. brevipes* 1 day old biofilm, G.mar_1d = *G. marina* 1 day old biofilm, G.mar_1w = *G. marina* 1 week old biofilm, G.mar_2w = *G. marina* 2 weeks old biofilm, G.mar_5w = *G. marina* 5 weeks old biofilm, Mix_3sp. = mix of three diatom species, Mix_6spS = mix of 6 microalgae species selected for positive effects on early settlement, Mix_6spG = mix of 6 microalgae species with positive effects on growth by 11 days, Mix_10sp. = mix of 10 microalgae species, nbc_1d = no biofilm control 1 day old, nbc_5w = no biofilm control 5 weeks old.

### Grammatophora marina biofilm

Having established that mixed microalgae biofilms do not have a synergistic effect on *Platynereis dumerilii* larval settlement compared to the most inductive single species biofilms, we next carried out settlement assays to further characterize the inductive properties of the most effective species of diatom biofilm, *Grammatophora marina*. We tested *G. marina* biofilms treated with heat or ethanol to reduce cell viability. We also tested *G. marina* filtrate and *G. marina* biofilm treated with antibiotics to separate any bacterial components from the diatom cell components. Antibiotic treatment of *G. marina* biofilm either before and after biofilm formation, or only after biofilm formation did not significantly alter the inductive capacity of the biofilm, while the filtrate of *G. marina*, which contained bacterial, but not diatom cells, did not induce significantly higher levels of *P. dumerilii* settlement than the no biofilm control (Fig. 4A, B). *G. marina* bifilms with reduced cell viability showed reduced levels of *P. dumerilii* settlement at 24 h and 48 h after induction. In particular, biofilm treated with EtOH or heated to 100°C induced levels of settlement similar to the no biofilm control. In these settlement assays, *G. marina* biofilm density, both of total cells and viable cells, showed a significant strong positive correlation with larval settlement rate according to Kendall’s Tau test (Supplementary data, Fig. S11). Ethanol, heat and antibiotic treatment of the *G. marina* biofilm all resulted in a small but significant reduction in death rate across the 48 h of the assays (Supplementary data, Fig. S10, Table S12).

**Figure 4.**
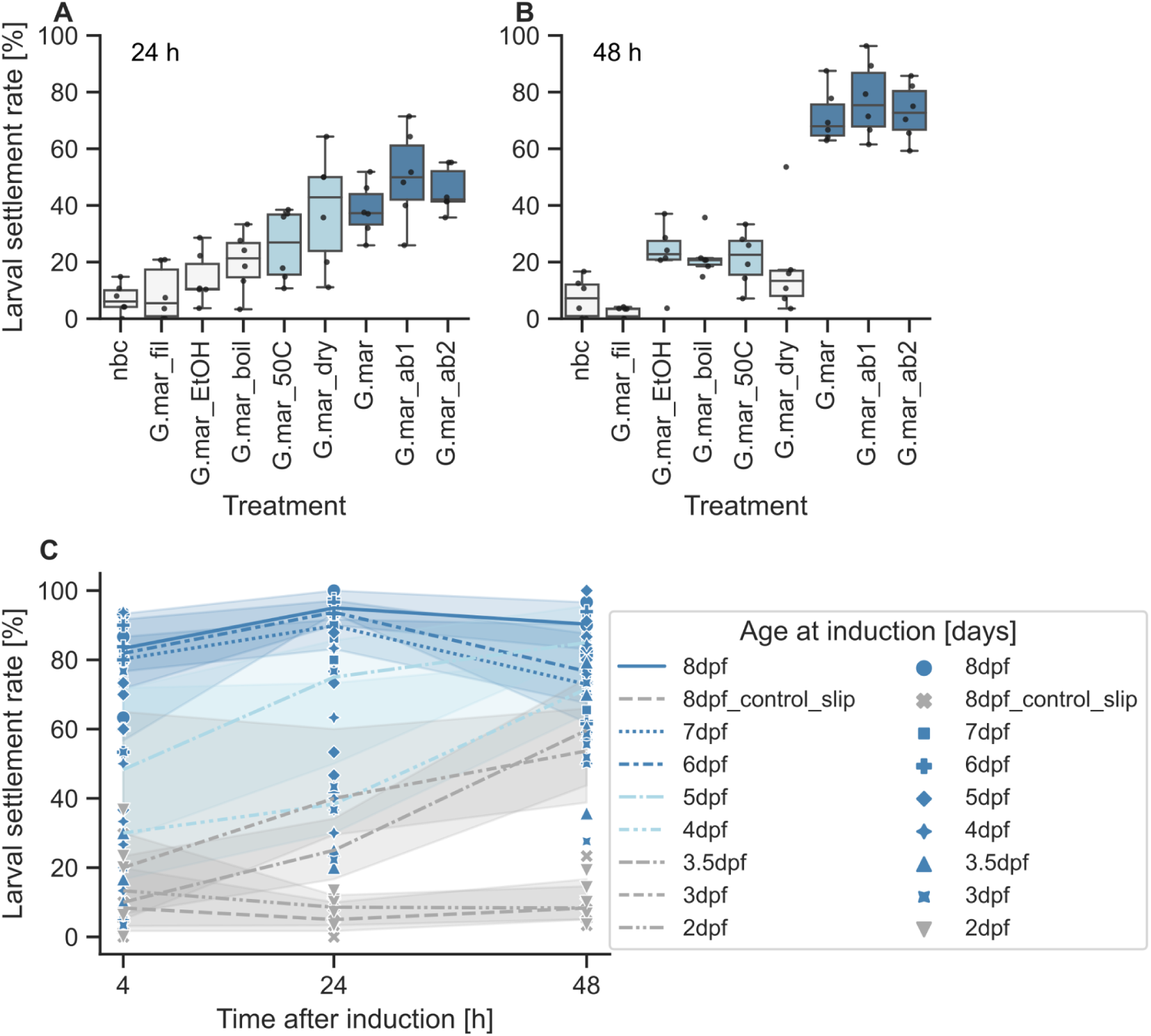
Characterization of *Platynereis dumerilii* larval settlement in response to *Grammatophora marina* biofilm. Box plots with scatter plot overlay of (A) % larval settlement in response to *G. marina* monospecies biofilms subjected to different treatments after 24 h and (B) 48 h exposure. Boxplots coloured according to the p-value result of a Mann-Whitney U rank test of % larval settlement at 24 and 48h, in each treatment versus the no biofilm control; p>0.05 = pale grey, p<0.05>0.00625 = light blue, p>0.00625 = dark blue. G.mar_fil = *G. marina* filtrate, G.mar_EtOH = EtOH-treated *G. marina*, G.mar_boil = boiled *G. marina*, G.mar_dry = *G. marina* 50°C overnight (dry), G.mar_50C = *G. marina* 50°C overnight (submerged), G.mar = 1 day old untreated *G. marina* biofilm, G.mar_ab1 = *G. marina* treated with antibiotics after biofilm formation, G.mar_ab2 = *G. marina* treated with antibiotics before and after biofilm formation, nbc = no biofilm control. (C) Line plot with scatter plot overlay of median % larval settlement over time for larvae introduced to *G. marina* biofilm at different initial ages (2-8 days old). Line shadows indicate 95% confidence interval. Lines are coloured according to the p-value result of a Mann-Whitney U rank test of % larval settlement at 4 h in each age group versus 8 days old larvae with a no biofilm control; p>0.05 = grey, p<0.05>0.00625 = light blue, p>0.00625 = dark blue. Raw data points coloured according to the p-value result of a Mann-Whitney U rank test of % larval settlement at 24 h in each age group versus 8 days old larvae with a no biofilm control; p>0.05 = grey, p<0.05>0.00625 = light blue, p>0.00625 = dark blue, (dpf = days post fertilization).

Settlement assays exposing *Platynereis dumerilii* larvae of different ages to one day old *Grammatophora marina* biofilm indicated that the capacity for larvae to respond to microalgal biofilms increases with age (Fig. 3C). Worms between four and eight days of age show significant increases in larval settlement rate compared to the no biofilm control even just 4 h after the start of the assay. In six to eight days old worms, between approximately 60 and 90% of worms had already settled on the *G. marina* biofilm within 4 h. In worms 3 days and older, significant levels of settlement were seen from 24 h after the start of the assay, but worms introduced to the assay at 2 days old larval stages never showed significantly increased rates of larval settlement across the 48 h of the assay. In the absence of a diatom biofilm, worms that were 8 days old at the start of the assay showed up to ∼20% settlement after 48 h, whereas 8 days old worms in the presence of *G. marina* biofilm reached a maximum of ∼96% settled after 48 h.

### Diatom effects on *Platynereis* growth

Similar to the positive effect of diatom species on early larval settlement, providing a diatom food source for *Platynereis dumerilii* following settlement also boosted growth of the worms over the first month of their life in our large scale culture (Fig. 5). Feeding young worms between 1 week and 1 month with a single diatom species (either *Phaeodactylum tricornutum* or *Skeletonema dohrnii*) increased size at 30 days by an additional ∼2 segments on average compared to feeding with a single green microalgal species (*Tetraselmis suecica* or *Nannochloropsis salina*). Highest growth after 30 days was seen in worms fed a mixed diet of either three microalgal species including one diatom species, or of five microalgal species including two diatom species, however increasing the species diversity in the mixed diet from three to five did not significantly change worm size. Median size achieved on mixed diets was 22-23 segments by 30 days. The largest worms recorded in these feeding trials had 37 segments. Worms of this size occurred under both mixed feeding regimes. Overall variance in size at 30 days under all diets was quite high, ranging from ∼ 14 segments in worms fed single species diets, to ∼26 segments in worms fed a mixed diet of five different microalgae species.

**Figure 5.**
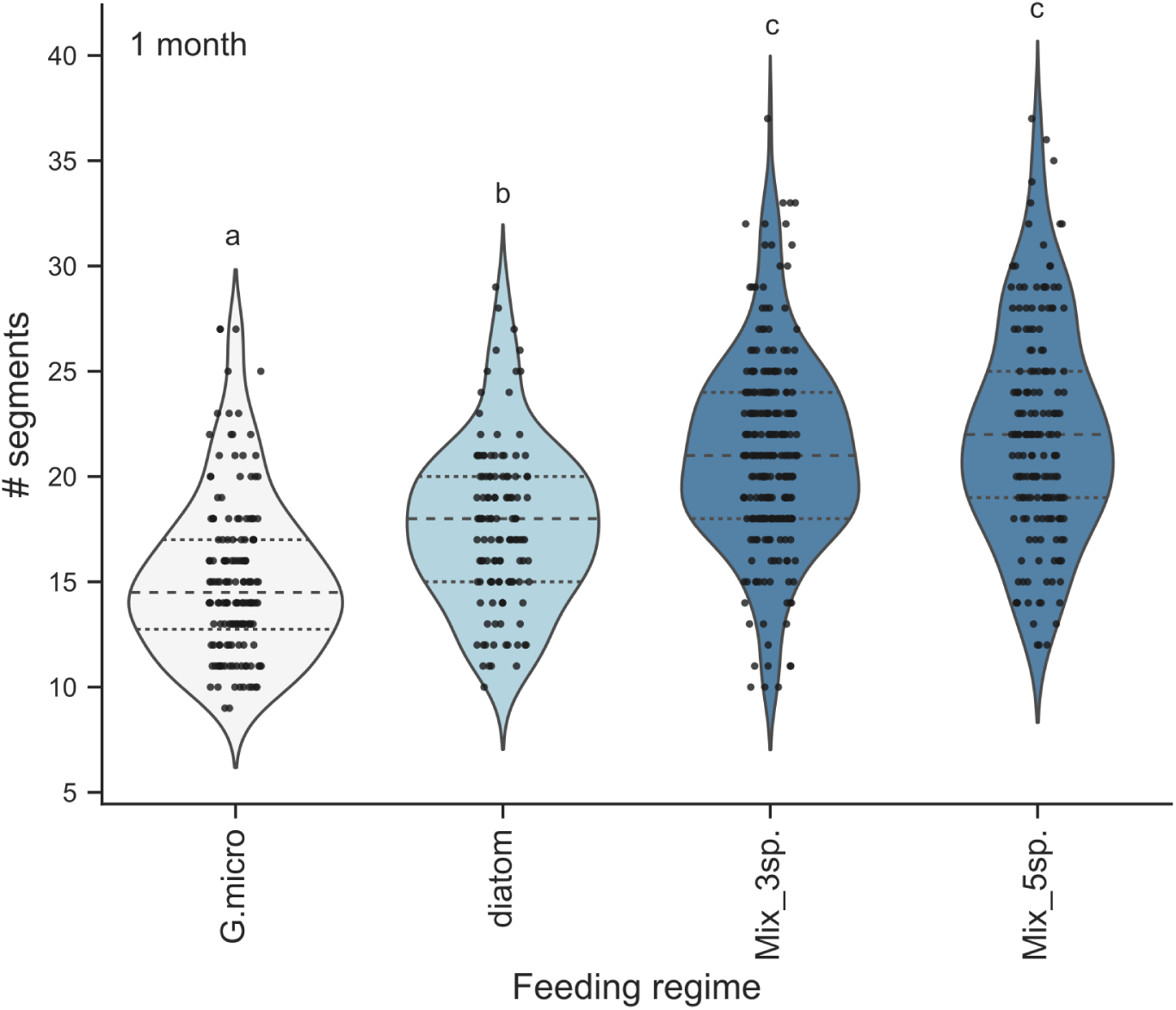
Effects of incorporation of diatoms into large scale *Platynereis dumerilii* culture. Violin plot with strip plot overlay of number of segments in individual 30 days old worms raised from 7 days on a single species green microalga (G.micro), a single species diatom (diatom), a mixed culture of two microalgae and one diatom species (Mix_3sp.), or a mixed culture of three microalgae and two diatoms (Mix_5sp.). Different feeding regimes are coloured based on their significance groups according to Kruskal-Wallis with Dunn’s post-hoc testing, which is also indicated above each group with ‘a’, ‘b’, or ‘c’. G.micro = green microalgae, Mix_3sp. = 2 microalgae plus 1 diatom species, Mix_5sp. = 3 microalgae plus 2 diatom species.

## 4. DISCUSSION

In this study we investigated a potential role for microbial biofilm in the induction of larval settlement of the marine worm *Platynereis dumerilii*. Through a series of behavioural assays, we established that microbial biofilm can induce an earlier onset of settlement and transition to a benthic lifestyle in the developing larva. The main component of biofilm that attracts larvae is microalgae, particularly diatoms, and *P. dumerilii* larvae respond to these microalgae in a species-specific manner.

### Microalgae as a larval settlement cue

Diatoms are already known to induce larval settlement in a species-specific manner in several other marine invertebrate larvae, although identification of species preferences is always limited by the number of species tested in each study. For example, mixed or monospecies diatom biofilms induced settlement and subsequent metamorphosis in the clam *Macoma balthica* (39), the polychaete *Hydroides elegans* (27,28,40), the bryozoan *Bugula neritina* (41,42), the slipper limpet *Crepidula onyx* (43), various abalone *Haliotis spp.* (44–47), the scallop *Argopecten purpuratus* (48), the acorn barnacle *Balanus amphitrite* (49,50), and the gastropod *Ilyanassa obsoleta* (51). Due to their positive effect on settlement, metamorphosis and subsequent juvenile growth, settlement plates coated with diatom-dominated biofilm are often implemented in shellfish and echinoderm aquaculture to enhance recruitment of spat (46,52–54). A growing body of evidence, including the results of this study, suggest that juvenile recruitment may be further optimized by selectively matching invertebrate larvae with their unique combination of preferred microalgal species.

As seems to be the case for other marine invertebrates whose larval settlement is induced by diatoms, *Platynereis dumerilii* larvae were particularly attracted to settle on biofilms dominated by benthic pennate diatom species including *Achnanthes* and *Nitzschia* species. Microalgae that induced higher levels of *P. dumerilii* settlement were not necessarily closely related to each other but the best inducers, including the coccolithophore *Chrysotila lamellosa*, share a common feature of secreting polysaccharide-rich extracellular polymeric substances (EPS) (55,56), leading us to conclude the particular element within the biofilm that attracts *P. dumerilii* may be found within the EPS. As EPS contains not only polysaccharides, but also proteins and lipids, further experimentation will be required to identify which specific aspects of EPS may be acting as infochemicals during *P. dumerilii* settlement. EPS was previously identified as being responsible for the inductive effect of diatom biofilms on larval settlement in *Hydroides elegans* (40), *Macoma balthica* (39) and *Balanus amphitrite* (49), with extracellular polysaccharides the most likely inductive cue.

Similarly, biofloc, which consists of extracellular polymers produced by diatoms and microbial protein, is recommended for culturing early life stages of the polychaete *Marphysa iloiloensis*, a species used in aquaculture, because it increases biomass in grow–out production (57).

Our settlement assays with *Grammatophora marina* biofilm treated with heat or ethanol to reduce viability indicate that the larval settlement activity of the biofilm is correlated with cell viability. This suggests that elements of the inductive settlement cue are derived from a microalgal metabolic pathway, and are transported to the EPS, where they are susceptible to degradation. A metabolically active biofilm is required to maintain inductive cue levels at a concentration high enough to be detected by the *P. dumerilii* larvae. A similar system has been proposed for the induction of *Hydroides elegans* larval settlement by bacterial biofilms (58).

We consistently recorded the highest levels of settlement in response to the chain-forming diatom species *Grammatophora marina*, although rates of larval settlement in response to this species were only marginally higher than those induced by *Achnanthes* and *Nitzschia* species. The fact that *G. marina* biofilm was equally effective in inducing initial *Platynereis dumerilii* larval settlement as various mixed species biofilms and that the effect was not significantly impacted by biofilm cell density or age, suggests that this species may produce a unique biochemical or physical signal to attract larvae during settlement. The highest levels of early settlement were induced by *G. marina*, but not the largest initial juvenile growth, indicating that the signal for a good place for initial settlement may be different from a general nutritive signal. A *Grammatophora* species co-occurs with a natural population of *P. dumerilii* in the Bay of Naples, Italy, in conjunction with a *Posidonia oceanica* seagrass bed habitat. A previous study of the feeding ecology of this *P. dumerilii* population found that *Grammatophora sp.* was one of the species found frequently in the benthic microalgal community of the seagrass and was also one of the most abundant genera of diatoms found in the faecal pellets of adult *P. dumerilii* collected from this habitat (23). Considering that the lab population of *P. dumerilii* used in this study was predominantly sourced originally from the Bay of Naples, *G. marina* may represent an ecologically relevant cue for *P. dumerilii* larval settlement, although we cannot confirm if the *Grammatophora sp*. described in Gambi *et al.* 2000 (23) was indeed *G. marina*. The physical structure of *G. marina*, as a branching diatom, may also contribute to its attractiveness to *P. dumerilii*. As young juveniles these worms prefer to feed on erect filamentous algae due to the shape and position of their jaws (22,23).

Results from our settlement assays indicate that unlike diatoms, bacteria in microbial biofilms are not involved in the regulation of *Platynereis dumerilii* larval settlement. Not only did monospecies bacterial biofilms fail to induce significant levels of larval settlement in *P. dumerilii*, but the antibiotic-treated *G. marina* diatoms were among the most effective for settlement induction. Antibiotic-treated *G. marina* was found to induce over 80% settlement after 48 h. In addition to this, filtrate of *G. marina*, which would still have retained any bacterial species present in the culture, also failed to induce settlement in *P. dumerilii*. This is surprising, as many previous studies have identified specific bacterial species from marine biofilm as inducers of larval settlement, including in other polychaete species (25,33). For example, *Pseudomonas marina* induced settlement and metamorphosis in the spirorbid polychaete *Janua brasiliensis* (59), and *Pseudoalteromonas luteoviolacea* (30,60,61), as well as *Cellulophaga lytica*, *Bacillus aquimaris* and *Staphylococcus warneri* (29,33,58), are all positive cues for metamorphosis in the polychaete *Hydroides elegans*. The Pseudoalteromonas genera in particular is a common cue for inducing larval settlement across multiple marine invertebrate phyla (62). Although we tested a *Pseudoalteromonas* species in our settlement assays with single-species bacterial biofilms, this did not induce significant *P. dumerilii* settlement over 48 h. We do not rule out that bacteria may also play a minor role in induction of *P. dumerilii* settlement, either directly or indirectly within a mixed biofilm - they can be important as initial colonizers of a surface to establish biofilm, or can contribute to biofilm robustness (63). In addition, there may still be a bacterial species that plays a major role in *P. dumerilii* settlement which we have not yet identified through our lab culture-based study. Our current findings indicate that diatoms are a more important cue than bacteria for *P. dumerilii* settlement, similar to larval settlement in some bryozoans, abalone, and barnacles (42,50,64). These species-specific differences in inductive cue specificity highlight the vast diversity that exists in the mechanisms that can trigger marine invertebrate larval settlement.

### Larval development and *Platynereis*-microalgae interactions

It is clear that in the case of *Platynereis dumerilii*, the cue for larval settlement is a signal that indicates the presence of food for the settled juvenile. Diatoms are an important dietary component for both juvenile and adult *Platynereis* species (23,65). Settlement assays with larvae of different ages indicate that *P. dumerilii* develop the initial capacity to respond to a *G. marina* biofilm, also known as larval competence, between three and four days old. This timing coincides with the development of the digestive tract, which is formed by the end of day four, while food intake usually begins between five and seven days of age (18,66,67). A distinguishing feature of *P. dumerilii* development between three and four days is the formation of additional sensory organs - tentacular cirri, which emerge from the lateral head and posterior, and antennal stubs, which grow on the anterior head (66). These sensory appendages may therefore be important for the detection of biofilms during settlement. Supporting this possibility, a study of the response of neuronal cells of different sensory head appendages in 6 days old *P. dumerilii* found that cells of the cirri, and especially the antennae, were responsive to glutamate, an amino acid, and sucrose, a sugar, indicative of a chemosensory role (68). If the antennae and cirri are already functional in younger larvae, it could be these which are responsible for microalgal biofilm cue detection during *P. dumerilii* larval settlement. Previous investigations of diverse polychaete species, including *Capitella teleta*, *Hydroides elegans*, *Phragmatopoma lapidosa* and *Phragmatopoma californica* (69–72), similarly indicate that the sensory cells responsible for selecting settlement sites may be located around the mouth, pharyngeal or head sensory appendages, rather than the larval apical sensory organ, which is already present in the earlier trochophore larval stage of development.

When two day old *Platynereis dumerilii* trochophore larvae are exposed to a *Grammatophora marina* biofilm, they fail to respond, and even show very low levels of larval settlement after exposure for 48 hours, by which time they are four days old and usually competent to settle. Similarly, premature exposure of some mollusc larvae to their preferred settlement cue can result in desensitization of the larvae and a lack of settlement response (73–76). This is thought to be due to habituation of the chemosensory receptors that detect the cue (77). This habituation effect provides a possible explanation for the lack of response of *P. dumerilii* to a microalgal biofilm when they encounter it prior to three days of age. The increased speed of response shown by *P. dumerilii* to biofilm as they age from three to eight days also suggests an accumulation of receptors sensitive to microalgal biofilm during development.

For *Platynereis dumerilii*, food is an essential requirement for growth and development beyond the three-segmented nectochaete larval stage. In the absence of food, larvae fail to add new posterior segments, and cannot complete the cephalic metamorphosis that occurs in 5-6 segmented juveniles (67). The nutritional profile of a microalga may therefore also be an important factor that determines its attractiveness to a *P. dumerilii* larva or recently settled juvenile. Different diatoms and other microalgal species can contain significantly different compositions of lipid, protein and sugars (78), which may act as a unique identifier to their consumers. Additionally, diatoms provide a primary source of omega-3 polyunsaturated fatty acids, particularly accumulating high levels of eicosapentaenoic acid (EPA) and docosahexaenoic acid (DHA) (79), which would be beneficial to a larva’s development and growth (80). This may explain why *P. dumerilii* fed a diet that includes diatoms undergo significantly more growth after one month compared to *P. dumerilii* fed a green microalgae alone. However, a mixed microalgal diet that includes both diatoms and other types of microalgae, such as green microalgae species, is optimal for *P. dumerilii*, as has been found for other animals (81), as this provides a more balanced nutrition and can further improve growth.

We found that levels of larval settlement in our initial settlement assays with *Tetraselmis suecica*, *Amphora coffeaeformis*, *Skeletonema dohrnii*, and *Phaeodactylum tricornutum* and *Chrysotila lamellosa* (Fig. 1C) did not always reflect results for the same microalgal species in subsequent settlement assays where we tested a broader range of species (Fig. 2). This may be due to changes over time in the health of the microalgal cultures, or a reflection of the variable genotypes of the *Platynereis dumerilii* larval batches used in different tests contributing to behavioural variation. Some diatoms, such as *P. tricornutum*, are plastic and able to change morphotype and lifestyle, switching between planktonic and benthic forms, which is also reflected in differing physiologies (82). Although we did not monitor individual cellular morphologies of different diatom species, this may have affected variability of their effectiveness in settlement assays. A previous study of *Achnanthes sp.* and *Nitzschia constricta* diatoms demonstrated that environmental conditions, including temperature and salinity, can impact diatom growth rate and the production of secreted extracellular polymers (EPS), which may be important as cues for invertebrate larvae (40,83). Species-specific preferences of microalgae in environmental culture conditions may also have impacted our results as we cultured all microalgal species with the same temperature and salinity. However, throughout both initial and subsequent settlement assays, the percentage of larval settlement remained consistently high for both *Grammatophora marina* and *Achnanthes brevipes* single species biofilms.

We also note that some diatoms are known for the secretion of noxious chemicals to deter their predators (84). Although some diatom species did not induce *Platynereis dumerilii* larval settlement at a significantly higher rate than the negative control, no species tested induced rates of settlement significantly lower than the negative control, suggesting that the larvae were not actively avoiding any of the tested species. Additionally, no microalgal species tested caused significantly higher rates of death in *P. dumerilii* larvae compared to the negative control, which would have indicated a strong toxic effect.

### Summary and outlook

We conclude that in larval settlement, *Platynereis dumerilii* is a generalist and will settle in the presence of any marine benthic biofilms that contain microalgae, particularly diatoms, which represent a future food source. However, the settlement decision is more robust and can occur earlier in development if the larvae receive signals from specific favoured species of diatom and coccolithophore, such as *Grammatophora marina*, some *Achnanthes* spp., *Nitzschia* spp., and the coccolithophore *Chrysotila lamellosa*, with preferred species being those known to produce EPS. This knowledge allows us to begin investigation of the link between the settlement-inducing environmental microalgal cues and the downstream internal molecular and nervous system signalling that governs settlement decisions in the *P. dumerilii* larva. The widespread occurrence of diatoms as a settlement cue for many invertebrate larvae, particularly those relevant to aquaculture, suggests that any findings on the signalling pathways of *P. dumerilii* settlement will also have relevance for these species.

In future, testing of EPS extracts of microalgal diatoms could confirm whether the settlement cue detected by *Platynereis dumerilii* larvae is indeed a component of the EPS, and help identify the molecular nature of the cue. Further investigation of larval-diatom interactions could be facilitated by the possibility of genome editing technologies in diatoms (85). This would allow more detailed investigations of the exact nature of the inductive molecules detected in diatoms by *P. dumerilii* larvae. For example, a specific metabolic pathway could be inhibited in a diatom biofilm which would then be tested in settlement assays to determine whether this altered the larval settlement activity.

As the incorporation of diatoms into *Platynereis dumerilii* culture can enhance settlement and subsequent growth, we recommend the inclusion of a select diatom species into their routine culture, and to commence administration of diatoms earlier than one week of age. This addition to *P. dumerilii* lab culture facility practice will help not only to enhance larval recruitment, but will contribute to reducing the overall generation time of *P. dumerilii*, an important factor for the efficient establishment of mutant lines in this research model.

## Acknowledgements

We thank Rebecca Turner, Sophie den Hartog, and Adam Johnstone for their contributions in running the *Platynereis* culture in the Marine Invertebrate Culture Unit (MICU) at the University of Exeter. Thanks also to Glen Wheeler and Katherine Helliwell, Marine Biological Association (MBA), Plymouth, UK for sharing their knowledge and expertise in microalgal biology and culture. Thanks to Stefano Migliorini for advice and suggestions on Python coding and statistical analyses.

## Author Contributions

Conceptualization: C.H., G.J., E.A.W.; Methodology: C.H., E.A.W.; Data analysis: C.H., E.A.W.; Investigation:C.H., E.A.W.; Writing - original draft: C.H, E.A.W; Writing - review & editing: C.H., G.J., E.A.W.; Project administration: E.A.W., C.H.; Funding Acquisition: E.A.W., G.J.

## Data Availability

Raw data files, scripts and macro for data analyses and figure generation are available on Github (https://github.com/MolMarSys-Lab/Platynereis-diatom_Hird_et_al_2024). Additional information and samples of raw images are available in the Supplementary Material.

