## Supplementary Material for "Microalgal biofilm induces larval settlement in the model marine worm *Platynereis dumerilii*"

**Supplementary Table S1.** BLAST results for bacterial species recovered from mixed biofilms grown in adult *Platynereis dumerilii* culture boxes. Species used for subsequent larval settlement assays are highlighted in green. File: 'Supplementary_Table_S1.xlsx'.

**Supplementary Table S2.** P-values of Mann-Whitney U Test on larval settlement in response to different bacteria and microalgae species (Fig. 1B), at 24h and 48h post induction.

| Species | 24h p-value | 48h p-value |
|---|---:|---:|
| No biofilm control | 1 | 1 |
| *Exiguobacterium sp. WR-24* | 0.549076 | 0.879389 |
| *Pseuduoalteromonas shioyasakiensis* | 0.051508 | 0.868661 |
| *Alteromonas sp. strain JLT1934* | 0.02558 | 0.535703 |
| *Halomonas venusta* | 0.101591 | 0.352897 |
| *Halobacillus sp. G-12* | 0.306492 | 0.172034 |
| *Tetraselmis suecica* | 0.040833 | 0.037188 |
| *Chrysotila lamellosa* | 0.029401 | 0.026519 |
| *Skeletonema dohrnii* | 0.02843 | 0.026519 |
| *Amphora coffeaeformis* | 0.029401 | 0.02558 |
| Mixed biofilm | 0.02843 | 0.02558 |
| *Phaeodactylum tricornutum* | 0.029401 | 0.02558 |

**Supplementary Table S3.** Estimated % coverage of different microalgae biofilms used in initial settlement assays (Fig. 1B). Based on single images of biofilmed coverslips taken at start of assay.

| Species | Estimated % coverage |
|---|---|
| *Amphora coffeaeformis* | 19.35% |
| *Chrysotila lamellosa* | 7.34% |
| Negative control | 0.007% |
| Positive control (mixed biofilm) | 92.29% |
| *Phaeodactylum tricornutum* | 6.685% |
| *Skeletonema dohrnii* | 13.952% |
| *Tetraselmis suecica* | 3.34% |

**Supplementary Table S4.** Metadata information for microalgae species tested in settlement and growth assays in Fig. 2. Table includes Species Name, Culture Collection ID, Family, Order, Group, Size (micron), Cell Shape, Call assemblage (chains vs solitary), and Culture Collection from which microalgae was sourced (Roscoff Culture Collection, France; Culture Collection of Algae and Protozoa, Scotland, UK; Marine Biological Association, Plymouth, UK; Belgian Coordinated Collections of Microorganisms/Diatom Collection Ghent, Belgium; Stazione Zoologica Napoli, Italy). File: 'Supplementary_Table_S4.xlsx'.

**Supplementary Table S5.** P-values of Mann-Whitney U Test on larval settlement in response to different microalgae species (Fig. 2), at 24h and 48h post induction.

| Species | 24h p-value | 48h p-value | Significance code (*) |
|---|---|---|---|
| Control_slip | 1 | 1 | |
| *Asterionellopsis_cf_glacialis* | 0.075669 | 0.917181 | |
| *Melosira_nummuloides* | 0.030626 | 0.406641 | * |
| *Staurosirella_pinnata* | 0.008661 | 0.37854 | * |
| *Navicula_phyllepta* | 0.536347 | 0.325845 | |
| *Amphora_sp_RCC7063* | 0.000283 | 0.276466 | ** |
| *Helicotheca_sp* | 0.060194 | 0.264673 | |
| *Coscinodiscus_sp* | 0.216341 | 0.20413 | |
| *Odontella_aurita* | 0.071481 | 0.196292 | |
| *Diploneis_sp* | 0.938334 | 0.196154 | |
| *Chaetoceros_convolutus* | 0.367113 | 0.08351 | |
| *Achnanthes_yaquinensis_0053* | 0.570838 | 0.082977 | |
| *Amphora_coffeaeformis* | 0.000819 | 0.074372 | ** |
| *Phaeodactylum_tricornutum* | 0.156533 | 0.070701 | |
| *Chaetoceros_cf_lauderi* | 0.979452 | 0.055556 | |
| *Conticribra_weisflogii* | 0.050324 | 0.043885 | * |
| *Entomoneis_alata* | 0.002598 | 0.04369 | ** |
| *Cyclotella_sp* | 0.180326 | 0.043592 | * |
| *Achnanthes_yaquinensis_0020* | 0.060155 | 0.028016 | * |
| *Chaetoceros_pseudocurvisetus* | 0.000513 | 0.027871 | *** |
| *Bacteriastrum_sp* | 0.000314 | 0.026361 | *** |
| *Gomphonema_sp* | 0.010016 | 0.016339 | ** |
| *Amphora_sp* | 0.004274 | 0.015127 | ** |
| *Hyalosira_sp* | 6.63E-05 | 0.011332 | *** |
| *Seminavis_robusta* | 0.11598 | 0.008397 | * |
| *Cylindrotheca_closterium* | 0.001999 | 0.006706 | ** |
| *Fragilariopsis_sp* | 0.15685 | 0.00574 | * |
| *Navicula_sp* | 0.000314 | 0.001264 | *** |
| *Chrysotila_lamellosa* | 0.000681 | 0.000952 | **** |
| *Surirella_sp* | 0.021927 | 0.000788 | *** |
| *Fragilariformia_virescens* | 0.000422 | 0.0006 | **** |
| *Tabularia_sp* | 0.02506 | 0.000494 | *** |
| *Skeletonema dohrnii* | 3.43E-05 | 0.000249 | **** |
| *Nitzschia_epithemoides* | 0.000233 | 0.000246 | **** |
| *Nitzschia_laevis* | 3.42E-05 | 0.000135 | **** |
| *Fragilaria_striatula* | 0.000113 | 0.000109 | **** |
| *Tetraselmis_suecica* | 0.06761 | 5.72E-05 | ** |
| *Achnanthes_sp* | 3.43E-05 | 3.29E-05 | **** |
| *Achnanthes_brevipes* | 3.43E-05 | 3.27E-05 | **** |

| | | | |
|---|---|---|---|
| *Nitzschia_ovalis* | 6.63E-05 | 3.29E-05 | **** |
| *Grammatophora_marina* | 3.43E-05 | 3.29E-05 | **** |

**Supplementary Figure S1.** Larval death during settlement assays in response to 40 species of microalgae (Fig. 2).

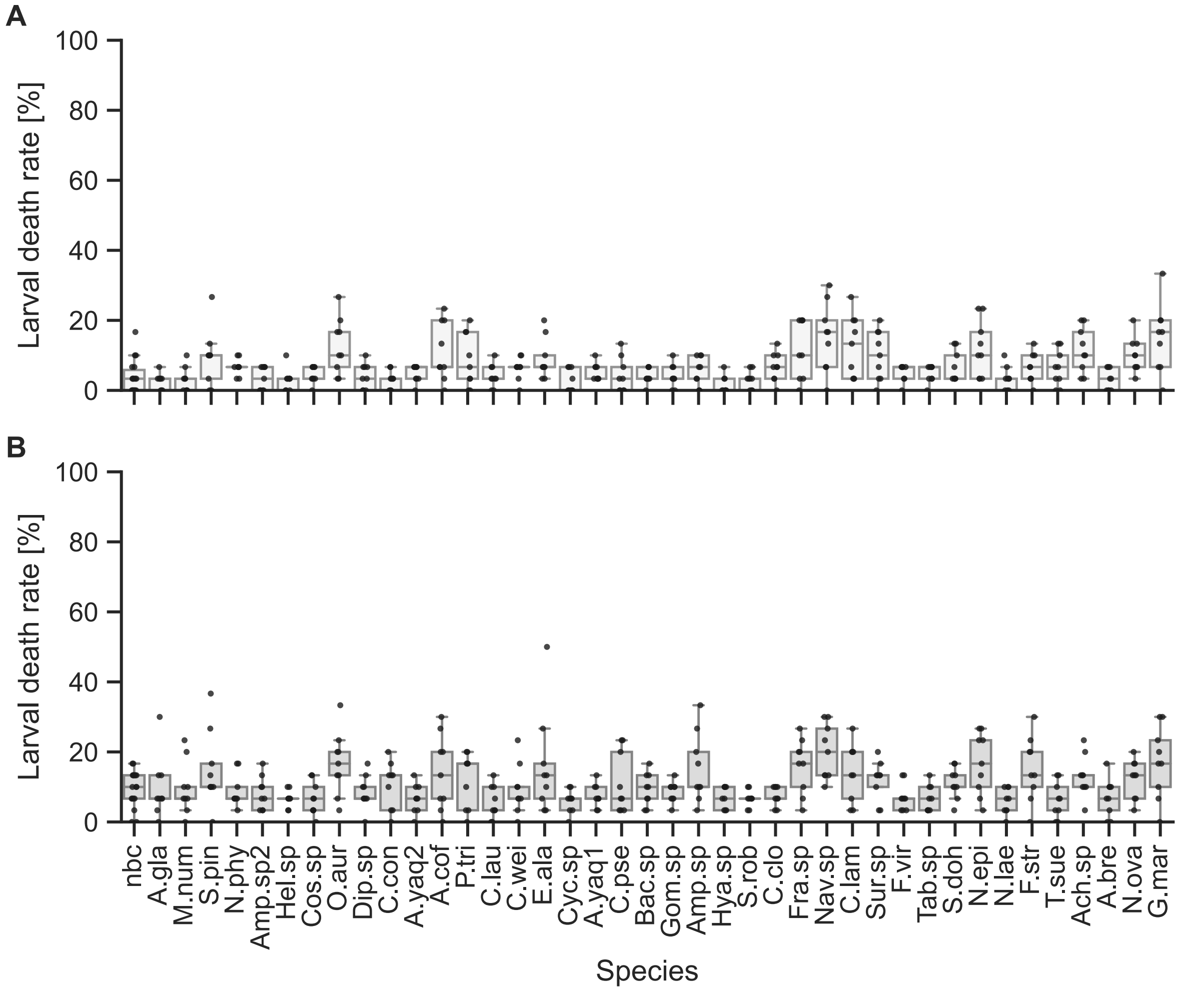


Fig. S1. Rates of death during settlement assays. Box plots with scatter plot overlay of (A) % larval death in response to different monospecies microalgal biofilms after 24h and (B) 48h exposure. A.gla = *Asterionellopsis cf glacialis*, M.num = *Melosira nummuloides*, S.pin = *Staurosirella pinnata*, N.phy = *Navicula phyllepta*, Amp.sp2 = *Amphora sp RCC7063*, Hel.sp = *Helicotheca sp.*, Cos.sp = *Coscinodiscus sp.*, O.aur = *Odontella aurita*, Dip.sp = *Diploneis sp.*, C.con = *Chaetoceros convolutus,* A.yaq2 = *Achnanthes yaquinensis DCG0053*, A.cof = *Amphora coffeaeformis*, P.tri = *Phaeodactylum tricornutum*, C.lau = *Chaetoceros cf lauderi*, C.wei = *Conticribra weisflogii*, E.ala = *Entomoneis alata*, Cyc.sp = *Cyclotella sp.*, A.yaq1 = *Achnanthes yaquinensis DCG0020*, C.pse = *Chaetoceros pseudocurvisetus*, Bac.sp = *Bacteriastrum sp.*, Gom.sp = *Gomphonema sp.,* Amp.sp = *Amphora sp.*, Hya.sp = *Hyalosira sp.*, S.rob = *Seminavis robusta*, C.clo = *Cylindrotheca closterium*, Fra.sp = *Fragilariopsis sp.*, Nav.sp = *Navicula sp.*, C.lam = *Chrysotila lamellosa*, Sur.sp = *Surirella sp.*, F.vir = *Fragilariformia virescens*, Tab.sp = *Tabularia sp.*, S.doh = *Skeletonema dohrnii*, N.epi = *Nitzschia epithemoides*, N.lae = *Nitzschia laevis*, F.str = *Fragilaria striatula*, T.sue = *Tetraselmis suecica*, Ach.sp = *Achnanthes sp.*, A.bre = *Achnanthes brevipes*, N.ova = *Nitzschia ovalis*, G.mar = *Grammatophora marina*, nbc = no biofilm control.

**Supplementary Table S6.** P-values of Mann-Whitney U Test on larval death in response to different microalgae species (Fig. 2), at 24h and 48h post induction.

| Species | 24h p-value | 48h p-value | Significance code (*) |
|---|---|---|---|
| Control_slip | 1 | 1 | |
| *Asterionellopsis_cf_glacialis* | 0.491712 | 0.912727 | |
| *Melosira_nummuloides* | 0.637352 | 0.869685 | |
| *Staurosirella_pinnata* | 0.306079 | 0.054584 | |
| *Navicula_phyllepta* | 0.578710 | 0.027455 | * |
| *Amphora_sp_RCC7063* | 0.401330 | 0.956546 | |
| *Helicotheca_sp* | 0.076275 | 1 | |
| *Coscinodiscus_sp* | 0.148010 | 0.270530 | |
| *Odontella_aurita* | 0.031817 | 0.002894 | ** |
| *Diploneis_sp* | 0.509203 | 0.608186 | |
| *Chaetoceros_convolutus* | 0.712354 | 0.978287 | |
| *Achnanthes_yaquinensis_0053* | 0.120887 | 0.198182 | |
| *Amphora_coffeaeformis* | 0.374366 | 0.014990 | * |
| *Phaeodactylum_tricornutum* | 0.297348 | 0.103588 | |
| *Chaetoceros_cf_lauderi* | 0.059346 | 0.500028 | |
| *Conticribra_weisflogii* | 0.600257 | 0.062296 | |
| *Entomoneis_alata* | 0.401554 | 0.012893 | * |
| *Cyclotella_sp* | 0.048394 | 0.659435 | * |
| *Achnanthes_yaquinensis_0020* | 0.178262 | 0.119251 | |
| *Chaetoceros_pseudocurvisetus* | 0.979185 | 0.891774 | |
| *Bacteriastrum_sp* | 0.772195 | 0.358448 | |
| *Gomphonema_sp* | 0.397939 | 0.500028 | |
| *Amphora_sp* | 0.240149 | 0.208444 | |
| *Hyalosira_sp* | 0.127498 | 0.228286 | |
| *Seminavis_robusta* | 0.076275 | 0.978287 | |
| *Cylindrotheca_closterium* | 0.162714 | 0.066794 | |
| *Fragilariopsis_sp* | 0.035001 | 0.011733 | ** |
| *Navicula_sp* | 0.003226 | 0.002908 | ** |
| *Chrysotila_lamellosa* | 0.165063 | 0.005512 | * |
| *Surirella_sp* | 0.313674 | 0.021195 | * |
| *Fragilariformia_virescens* | 0.152874 | 0.140374 | |
| *Tabularia_sp* | 0.109244 | 0.198182 | |
| *Skeletonema dohrnii* | 0.544768 | 0.060714 | |
| *Nitzschia_epithemoides* | 0.056795 | 0.005467 | * |
| *Nitzschia_laevis* | 0.055784 | 0.891811 | |
| *Fragilaria_striatula* | 0.104270 | 0.098546 | |
| *Tetraselmis_suecica* | 0.127134 | 0.173344 | |
| *Achnanthes_sp* | 0.256235 | 0.003163 | * |
| *Achnanthes_brevipes* | 0.218145 | 0.934792 | |
| *Nitzschia_ovalis* | 0.245114 | 0.003780 | * |
| *Grammatophora_marina* | 0.031807 | 0.002894 | ** |

**Supplementary Figure S2.** Biofilm % coverage for the 39 species/40 strains of microalgae used in settlement assays.

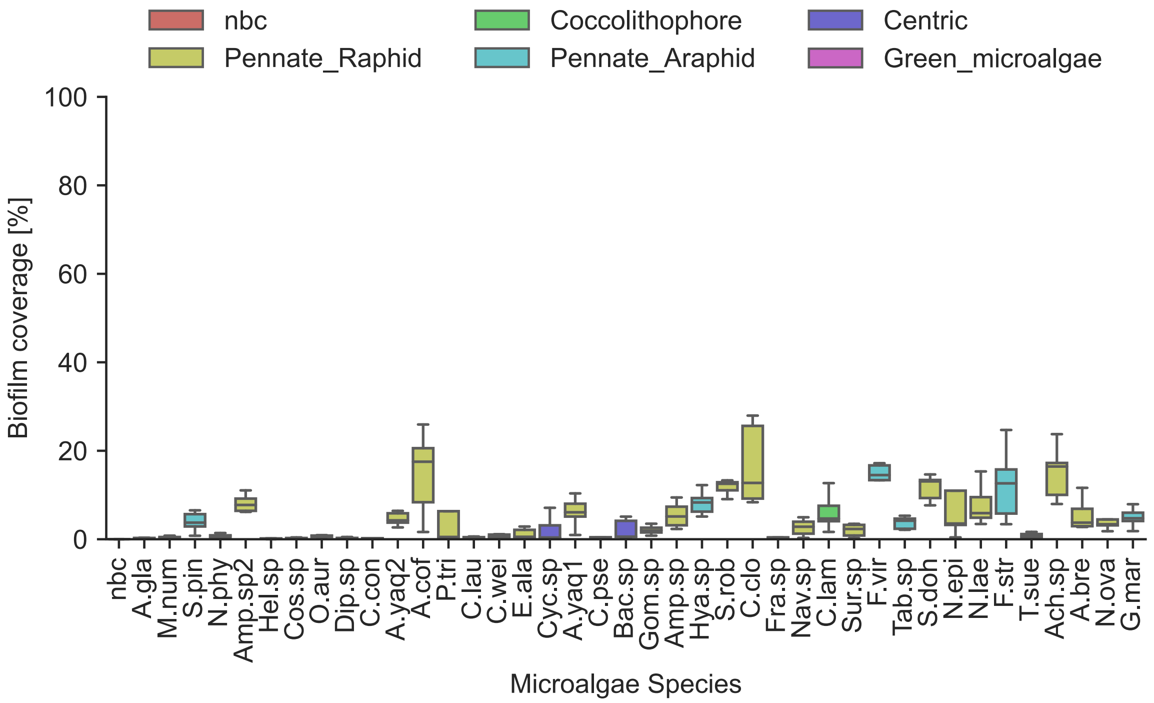


Fig. S2. Percent biofilm coverage during settlement assays. Box plots of biofilm coverage [%] for different monospecies microalgal biofilms at start of assay. A.gla = *Asterionellopsis cf glacialis*, M.num = *Melosira nummuloides*, S.pin = *Staurosirella pinnata*, N.phy = *Navicula phyllepta*, Amp.sp2 = *Amphora sp RCC7063*, Hel.sp = *Helicotheca sp.*, Cos.sp = *Coscinodiscus sp.*, O.aur = *Odontella aurita*, Dip.sp = *Diploneis sp.*, C.con = *Chaetoceros convolutus,* A.yaq2 = *Achnanthes yaquinensis DCG0053*, A.cof = *Amphora coffeaeformis*, P.tri = *Phaeodactylum tricornutum*, C.lau = *Chaetoceros cf lauderi*, C.wei = *Conticribra weisflogii*, E.ala = *Entomoneis alata*, Cyc.sp = *Cyclotella sp.*, A.yaq1 = *Achnanthes yaquinensis DCG0020*, C.pse = *Chaetoceros pseudocurvisetus*, Bac.sp = *Bacteriastrum sp.*, Gom.sp = *Gomphonema sp.,* Amp.sp = *Amphora sp.*, Hya.sp = *Hyalosira sp.*, S.rob = *Seminavis robusta*, C.clo = *Cylindrotheca closterium*, Fra.sp = *Fragilariopsis sp.*, Nav.sp = *Navicula sp.*, C.lam = *Chrysotila lamellosa*, Sur.sp = *Surirella sp.*, F.vir = *Fragilariformia virescens*, Tab.sp = *Tabularia sp.*, S.doh = *Skeletonema dohrnii*, N.epi = *Nitzschia epithemoides*, N.lae = *Nitzschia laevis*, F.str = *Fragilaria striatula*, T.sue = *Tetraselmis suecica*, Ach.sp = *Achnanthes sp.*, A.bre = *Achnanthes brevipes*, N.ova = *Nitzschia ovalis*, G.mar = *Grammatophora marina*, nbc = no biofilm control (see also Fig. S4 for example images).

Kendall's Tau correlation test results, for correlation between biofilm density and larval settlement rate:

(1) All 40 strains tested, settlement at 24 hours post induction (hpi)
Significance Result: statistic = 0.257, p-value = 6.36E-13
(2) All 40 strains tested, settlement at 48 hours post induction (hpi)
Significance Result: statistic = 0.305, p-value = 8.15E-18
(3) For 15 significantly inductive species at 24hpi
Significance Result: statistic = 0.099, p-value = 0.09

(4) For 15 significantly inductive species at 48hpi
Significance Result: statistic = 0.142, p-value = 0.015

**Supplementary Figure S3.** Plots of % larval settlement at 24hpi in response to 40 different monospecies biofilms (Fig. 2), grouped by 'Group', 'Size Class', Cell Shape', and 'Cell Assembly'.

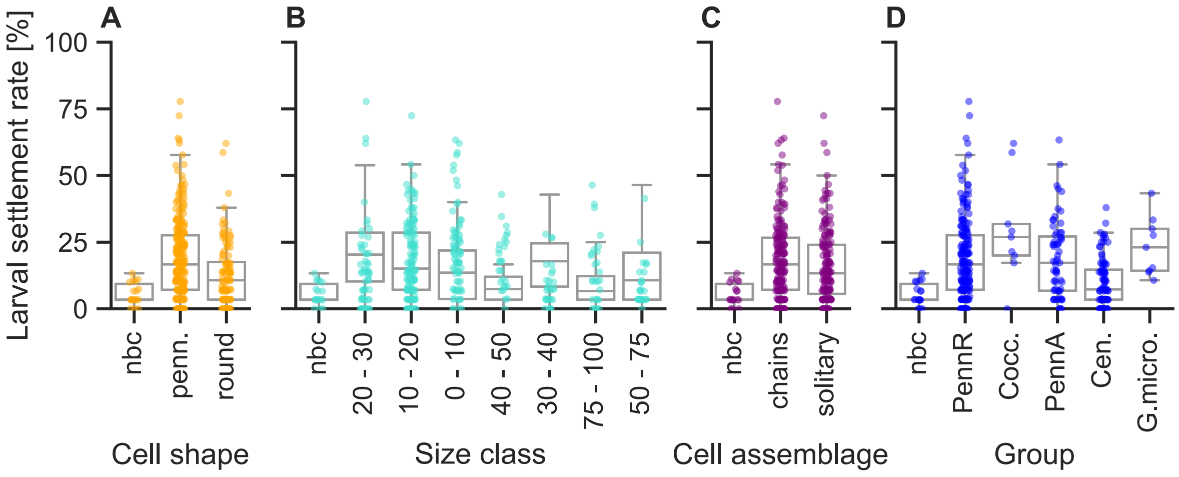


Fig.S3. Box plots with scatter plot overlay of % larval settlement at 24hpi in response to 40 different monospecies biofilms, grouped by (A) Cell Shape, (B) Size class (C) Cell assemblage, and (D) Group. 'Group'. nbc = no biofilm control, penn. = pennate, PennR = pennate raphid, Cocc. = coccolithophore, PennA = pennate araphid, Cen. = centric, G.micro. = green microalgae.

Corresponding Kruskal-Wallis Test with Dunn's posthoc testing values:

(1) Cell shape (p-value cut-off = 0.025)

KruskalResult (statistic=32.03373719598774, pvalue=1.1065278536725637e-07)

| | nbc | pennate | round |
|---|---|---|---|
| nbc | 1.000000 | 0.000012 | 0.023822 |
| pennate | 0.000012 | 1.000000 | 0.000285 |
| round | 0.023822 | 0.000285 | 1.000000 |

(2) Size class (p-value cut-off = 0.007)

KruskalResult (statistic=51.87448854174704, pvalue=6.180718899398846e-09)

| | nbc | 0-10 | 10-20 | 20-30 | 30-40 | 40-50 | 50-75 |
|---|---|---|---|---|---|---|---|
| nbc | 1.000000 | 0.000016 | 0.000141 | 0.056552 | 1.000000 | 0.004742 | 1.000000 |
| 0-10 | 0.000016 | 1.000000 | 1.000000 | 0.221623 | 0.001143 | 1.000000 | 0.000497 |
| 10-20 | 0.000141 | 1.000000 | 1.000000 | 1.000000 | 0.009949 | 1.000000 | 0.004535 |
| 20-30 | 0.056552 | 0.221623 | 1.000000 | 1.000000 | 1.000000 | 1.000000 | 1.000000 |
| 30-40 | 1.000000 | 0.001143 | 0.009949 | 1.000000 | 1.000000 | 0.160543 | 1.000000 |
| 40-50 | 0.004742 | 1.000000 | 1.000000 | 1.000000 | 0.160543 | 1.000000 | 0.098306 |
| 50-75 | 1.000000 | 0.000497 | 0.004535 | 1.000000 | 1.000000 | 0.098306 | 1.000000 |
| 75-100 | 0.378464 | 0.336537 | 1.000000 | 1.000000 | 1.000000 | 1.000000 | 1.000000 |

```
       75-100
nbc    0.378464
0-10   0.336537
10-20  1.000000
20-30  1.000000
30-40  1.000000
40-50  1.000000
50-75  1.000000
75-100 1.000000
```

(3) Cell assemblage (p-value cut-off = 0.025)

KruskalResult (statistic=18.99181096268572, pvalue=7.515894040751246e-05)

```
          nbc       chains    solitary
nbc       1.000000  0.000012  0.023822
chains    0.000012  1.000000  0.000285
solitary  0.023822  0.000285  1.000000
```

(5) Group (p-value cut-off = 0.01)

KruskalResult (statistic=52.72037415187003, pvalue=3.838606646867595e-10)

```
         nbc       PennR     Cocc      PennA     Cen       G.micro
nbc      1.000000  0.000076  0.000495  0.000393  0.943349  0.001173
PennR    0.000076  1.000000  1.000000  1.000000  0.000007  1.000000
Cocc     0.000495  1.000000  1.000000  1.000000  0.007677  1.000000
PennA    0.000393  1.000000  1.000000  1.000000  0.001348  1.000000
Cen      0.943349  0.000007  0.007677  0.001348  1.000000  0.018010
G.micro  0.001173  1.000000  1.000000  1.000000  0.018010  1.000000
```

**Supplementary Table S7.** P-values of Mann-Whitney U Test on *Platynereis* length at 11 days in response to different microalgae species (Fig. 2C).

| Species | 24h p-value | Significance code (*) |
|---|---|---|
| Control_slip | 1 | |
| *Asterionellopsis_cf_glacialis* | 0 | ** |
| *Melosira_nummuloides* | 0 | ** |
| *Staurosirella_pinnata* | 0.554623 | |
| *Navicula_phyllepta* | 0.015522 | * |
| *Amphora_sp_RCC7063* | 0 | ** |
| *Helicotheca_sp* | 0.000019 | ** |
| *Coscinodiscus_sp* | 0.000002 | ** |
| *Odontella_aurita* | 0.493261 | |
| *Diploneis_sp* | 0.000003 | ** |
| *Chaetoceros_convolutus* | 0 | ** |
| *Achnanthes_yaquinensis_0053* | 0 | ** |
| *Amphora_coffeaeformis* | 0 | ** |
| *Phaeodactylum_tricornutum* | 0.836660 | |
| *Chaetoceros_cf_lauderi* | 0.019835 | * |
| *Conticribra_weisflogii* | 0.798572 | |

| | | |
|---|---|---|
| *Entomoneis_alata* | 0.008189 | * |
| *Cyclotella_sp* | 0 | ** |
| *Achnanthes_yaquinensis_0020* | 0 | ** |
| *Chaetoceros_pseudocurvisetus* | 0 | ** |
| *Bacteriastrum_sp* | 0 | ** |
| *Gomphonema_sp* | 0 | ** |
| *Amphora_sp* | 0 | ** |
| *Hyalosira_sp* | 0 | ** |
| *Seminavis_robusta* | 0 | ** |
| *Cylindrotheca_closterium* | 0.000012 | ** |
| *Fragilariopsis_sp* | 0.032284 | * |
| *Navicula_sp* | 0.125416 | |
| *Chrysotila_lamellosa* | 0 | ** |
| *Surirella_sp* | 0.791783 | |
| *Fragilariformia_virescens* | 0 | ** |
| *Tabularia_sp* | 0 | ** |
| *Skeletonema dohrnii* | 0 | ** |
| *Nitzschia_epithemoides* | 0.318205 | |
| *Nitzschia_laevis* | 0 | ** |
| *Fragilaria_striatula* | 0 | ** |
| *Tetraselmis_suecica* | 0.057497 | |
| *Achnanthes_sp* | 0 | ** |
| *Achnanthes_brevipes* | 0 | ** |
| *Nitzschia_ovalis* | 0 | ** |
| *Grammatophora_marina* | 0 | ** |

**Supplementary Figure S4.** Example light micrograph images for 40 different monospecies biofilms used in settlement assays (Fig. 2)

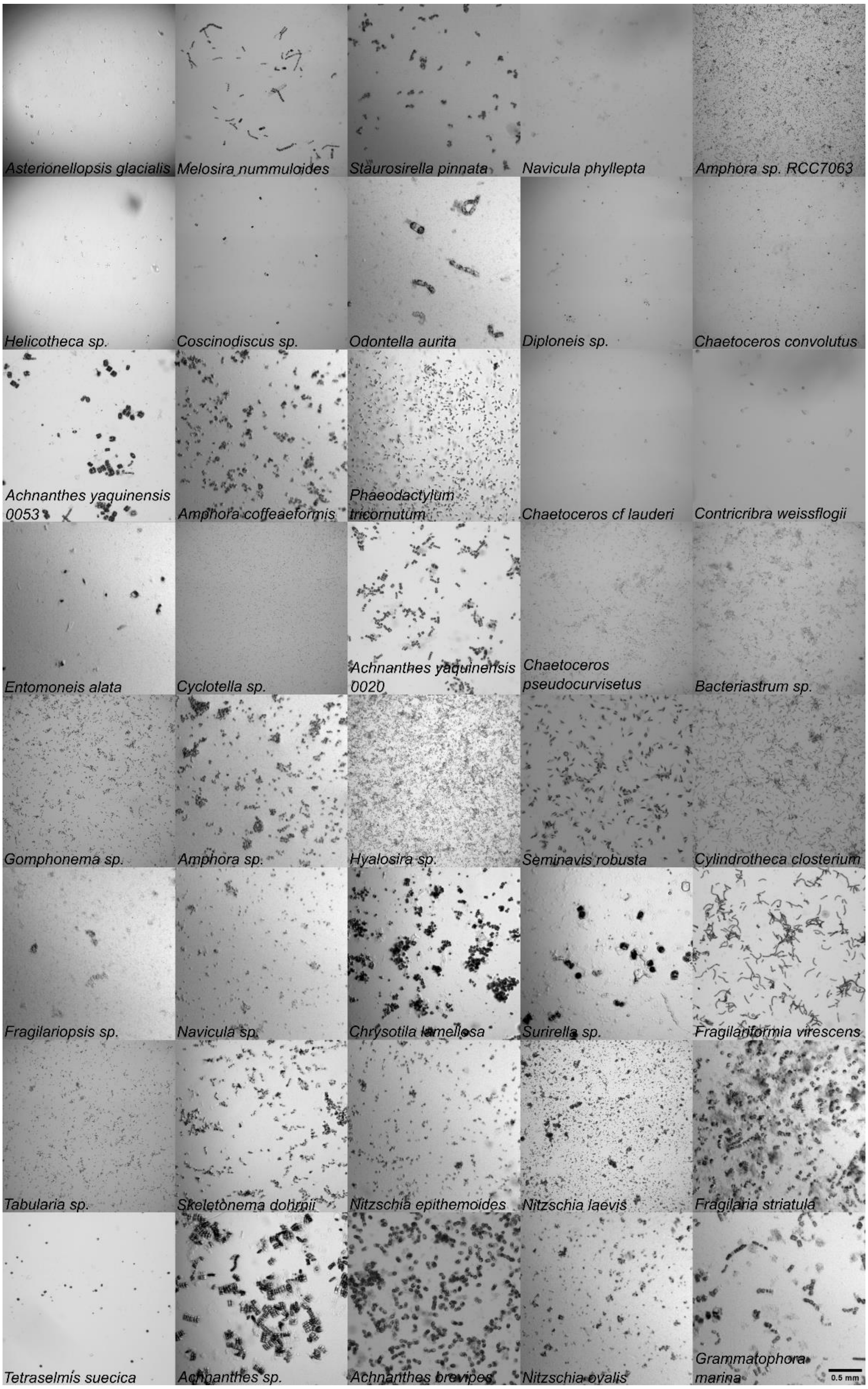


Fig. S4. Example light micrograph images of biofilms used in single-species settlement assays. Images ordered according to % larval settlement at 24 h (lowest to highest). Species name in bottom left corner. Scale bar 0.5mm.

**Supplementary Figure S5.** Example light micrograph images for postlarval growth at 11 days in response to 40 different monospecies biofilms used in settlement assays (Fig. 2C).

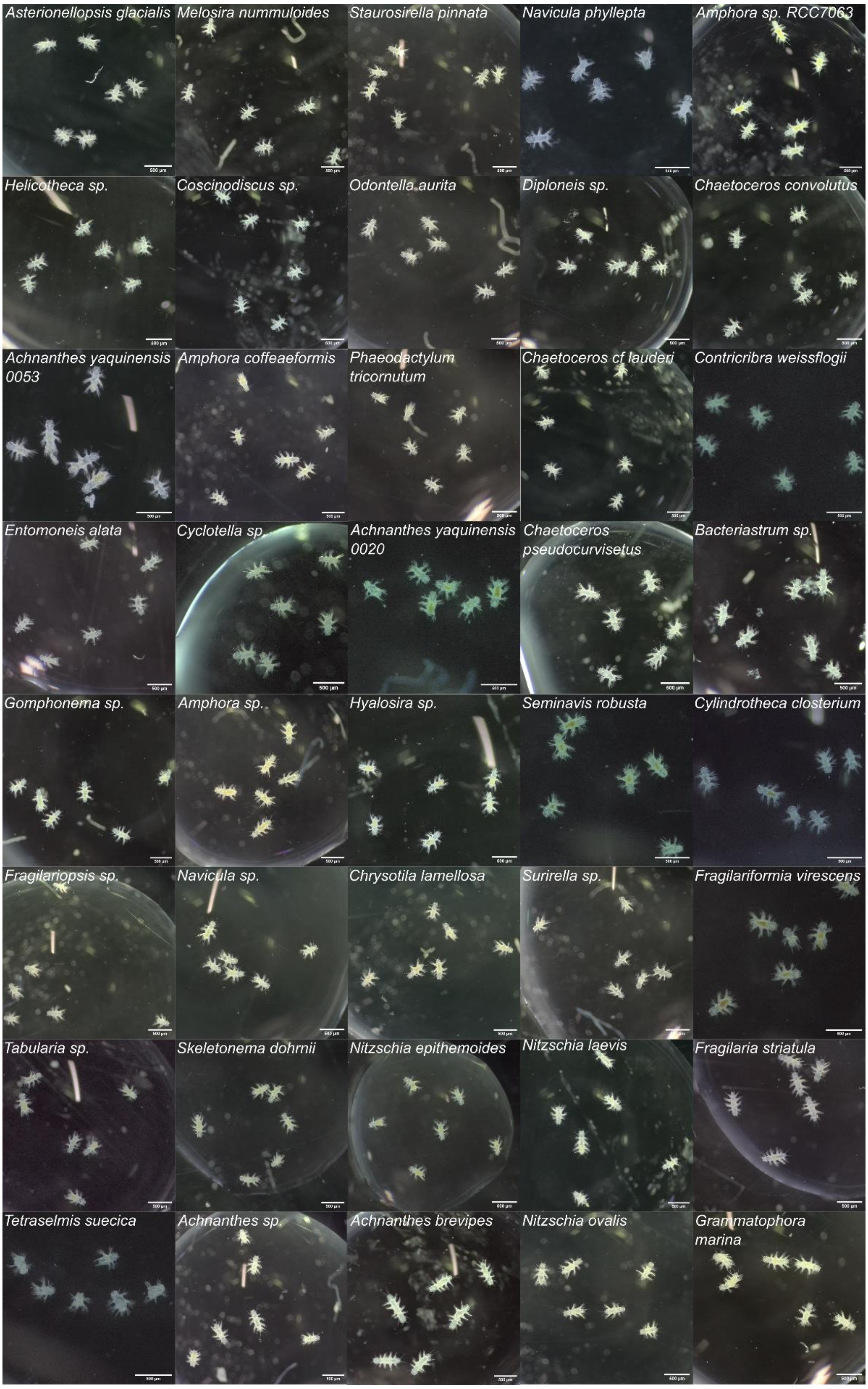


Fig. S5. Example light micrograph images of 11 days old postlarvae used in single-species settlement assays. Biofilm species name in top left corner. Scale bar 500 μm.

**Supplementary Table S8.** P-values of Mann-Whitney U Test on larval settlement in response to different microalgae species or mixtures (Fig. 3), at 24h and 48h post induction.

| Biofilm composition | 24h p-value | 48h p-value | Significance code (*) |
|---|---|---|---|
| NEG_CNTRL_1day | 1 | 1 | |
| NEG_CNTRL_5weeks | 0.048885 | 0.436661 | * |
| 3_species_mix | 0.004922 | 0.025974 | *** |
| Grammatophora_marina_1week | 0.027532 | 0.02381 | ** |
| Grammatophora_marina_2weeks | 0.027532 | 0.02381 | ** |
| Grammatophora_marina_5weeks | 0.027532 | 0.02381 | ** |
| Achnanthes_brevipes_1day | 0.004922 | 0.010272 | *** |
| Grammatophora_marina_1day | 0.004998 | 0.008658 | *** |
| 10_species_mix | 0.004998 | 0.004998 | **** |
| 6_species_mix_settle | 0.004998 | 0.002165 | **** |
| 6_species_mix_settle_grow | 0.004998 | 0.002165 | **** |

**Supplementary Figure S6.** Larval death during settlement assays in response to different microalgae species or mixtures (Fig. 3), at 24h and 48h post induction.

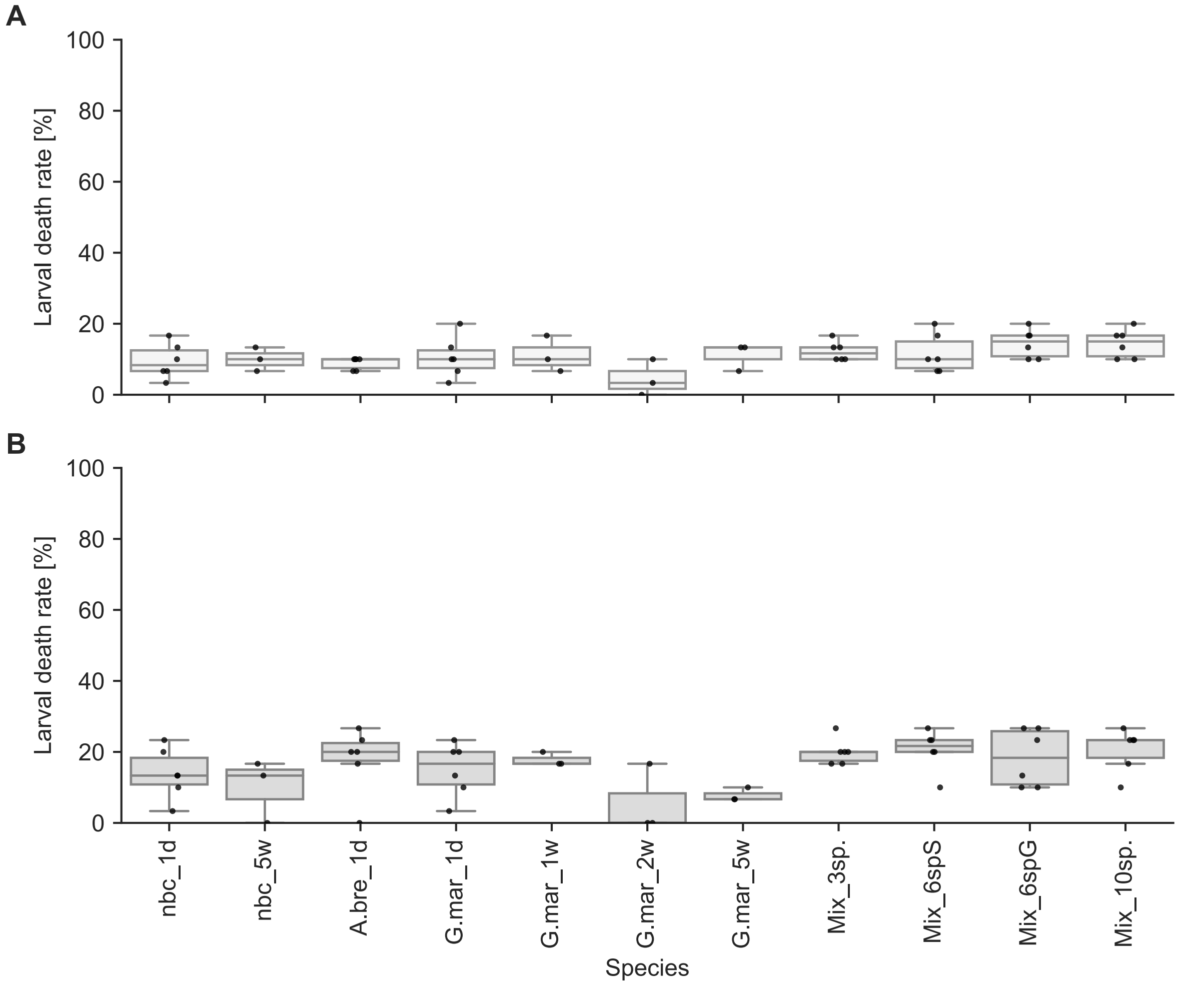


Fig. S6. Rates of death during settlement assays. Box plots with scatter plot overlay of (A) % larval death in response to different mono- or mixed-species microalgal biofilms after 24h and (B) 48h exposure. A.bre_1d= *Achnanthes brevipes* 1 day old, G. mar_1d =

*Grammatophora marina* 1 day old, G.mar_1w = *G. marina* 1 week old, G.mar_2w = *G. marina* 2 weeks old, G.mar_5w = *G. marina* 5 weeks old, Mix_3sp. = mixture of 3 microalgae species, Mix_spS = mixture of 6 species optimized for settlement, Mix_6spG = mixture of 6 species optimized for growth, Mix_10sp. = mixture of 10 microalgae species, nbc_1d = 1 day old no biofilm control, nbc_5w = 5 weeks old no biofilm control.

**Supplementary Table S9.** P-values of Mann-Whitney U Test on larval death in response to different mono- or mixed microalgae species biofilms (Fig. 3), at 24h and 48h post induction.

| Biofilm composition | 24h p-value | 48h p-value | Significance code (*) |
|---|---|---|---|
| NEG_CNTRL_1day | 1.000000 | 1.000000 | |
| NEG_CNTRL_5weeks | 0.693641 | 0.894626 | |
| 3_species_mix | 0.140920 | 0.284067 | |
| Grammatophora_marina_1week | 0.432768 | 0.691102 | |
| Grammatophora_marina_2weeks | 0.241317 | 0.239317 | |
| Grammatophora_marina_5weeks | 0.191063 | 0.688500 | |
| Achnanthes_brevipes_1day | 0.331544 | 1.000000 | |
| Grammatophora_marina_1day | 0.870283 | 0.806840 | |
| 10_species_mix | 0.120143 | 0.103095 | |
| 6_species_mix_settle | 0.142368 | 0.510538 | |
| 6_species_mix_settle_grow | 0.415063 | 0.103095 | |

**Supplementary Figure S7.** Biofilm % coverage for the different mono- and mixed-species microalgae biofilms used in settlement assays (Fig. 3)

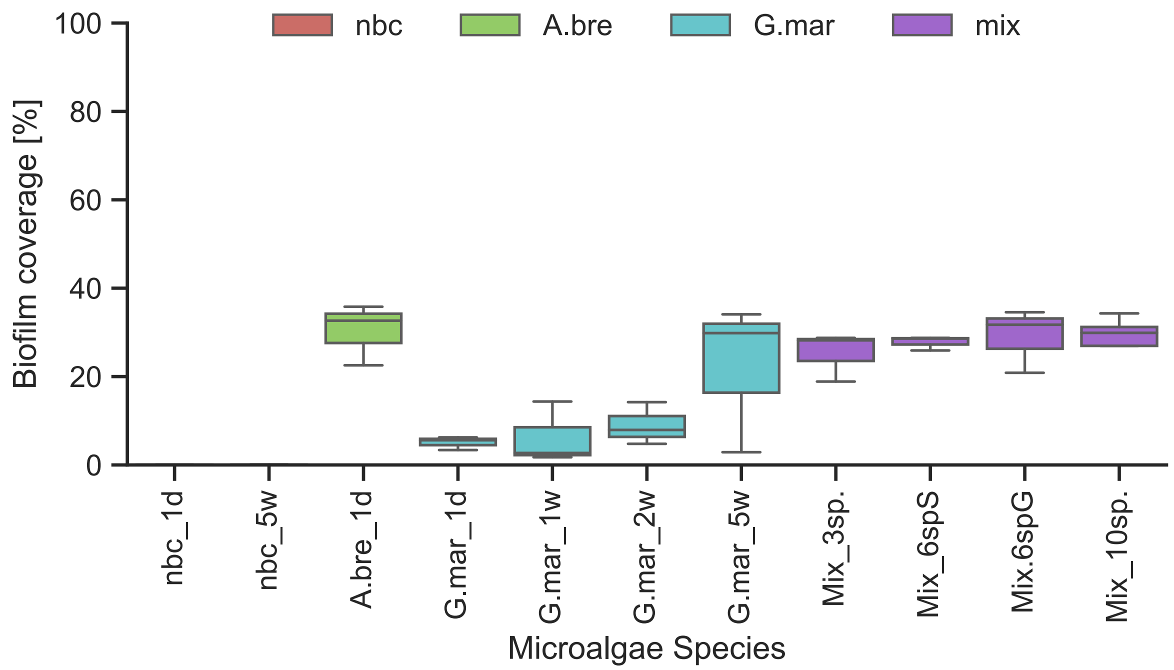


Fig. S7. Percent biofilm coverage during settlement on single or mixed microalgae species biofilms. Box plots of biofilm coverage [%] for different monospecies microalgal biofilms at start of assay. A.bre_1d= *Achnanthes brevipes* 1 day old, G. mar_1d = *Grammatophora marina* 1 day old, G.mar_1w = *G. marina* 1 week old, G.mar_2w = *G. marina* 2 weeks old,

G.mar_5w = *G. marina* 5 weeks old, Mix_3sp. = mixture of 3 microalgae species, Mix_spS = mixture of 6 species optimized for settlement, Mix_6spG = mixture of 6 species optimized for growth, Mix_10sp. = mixture of 10 microalgae species, nbc_1d = 1 day old no biofilm control, nbc_5w = 5 weeks old no biofilm control (see also Fig. S8 for example images)

**Kendall's Tau correlation test results**, for correlation between biofilm density and larval settlement rate:

(1) All 9 biofilms tested, settlement at 24 hours post induction (hpi)
Significance Result: statistic = -0.056, p-value = 0.919455
(2) All 9 biofilms tested, settlement at 48 hours post induction (hpi)
Significance Result: statistic = 0.278, p-value = 0.358488

Corresponding Kruskal-Wallis Test with Dunn's posthoc testing values:

(1) *G. marina* biofilms of different age, settlement at 24 hpf (p-value cut-off = 0.05)

KruskalResult (`statistic=15.678783921056462, pvalue=0.007823584368925156`)

| | nbc_1d | nbc_5w | G.mar_1d | G.mar_1w | G.mar_2w | G.mar_5w |
|---|---|---|---|---|---|---|
| nbc_1d | 1.000000 | 1.000000 | 0.410781 | 1.000000 | 0.017182 | 0.047370 |
| nbc_5w | 1.000000 | 1.000000 | 1.000000 | 1.000000 | 0.488250 | 0.906511 |
| G.mar_1d | 0.410781 | 1.000000 | 1.000000 | 1.000000 | 1.000000 | 1.000000 |
| G.mar_1w | 1.000000 | 1.000000 | 1.000000 | 1.000000 | 1.000000 | 1.000000 |
| G.mar_2w | 0.017182 | 0.488250 | 1.000000 | 1.000000 | 1.000000 | 1.000000 |
| G.mar_5w | 0.047370 | 0.906511 | 1.000000 | 1.000000 | 1.000000 | 1.000000 |

(2) Single vs mixed 1 day old biofilms, settlement at 24 hpf (p-value cut-off = 0.05)

KruskalResult (`statistic=18.74024340770792, pvalue=0.004625571161867118`)

| | nbc_1d | A.bre_1d | G.mar_1d | Mix_3sp. | Mix_6spS | Mix_6spG |
|---|---|---|---|---|---|---|
| 7 | | | | | | |
| nbc_1d | 1.000000 | 0.758431 | 0.137258 | 1.000000 | 0.021438 | 0.012864 |
| A.bre_1d | 0.758431 | 1.000000 | 1.000000 | 1.000000 | 1.000000 | 1.000000 |
| G.mar_1d | 0.137258 | 1.000000 | 1.000000 | 1.000000 | 1.000000 | 1.000000 |
| Mix_3sp. | 1.000000 | 1.000000 | 1.000000 | 1.000000 | 1.000000 | 1.000000 |
| Mix_6spS | 0.021438 | 1.000000 | 1.000000 | 1.000000 | 1.000000 | 1.000000 |
| Mix_6spG | 0.012864 | 1.000000 | 1.000000 | 1.000000 | 1.000000 | 1.000000 |
| Mix_10sp. | 0.014025 | 1.000000 | 1.000000 | 1.000000 | 1.000000 | 1.000000 |

| | Mix_10sp. |
|---|---|
| nbc_1d | 0.014025 |
| A.bre_1d | 1.000000 |
| G.mar_1d | 1.000000 |
| Mix_3sp. | 1.000000 |
| Mix_6spS | 1.000000 |
| Mix_6spG | 1.000000 |
| Mix_10sp. | 1.000000 |

**Supplementary Table S10.** P-values of Mann-Whitney U Test on *Platynereis* length at 11 days in response to different microalgae species (Fig. 2C).

| Biofilm composition | 11 days p-value | Significance code (*) |
|---|---|---|
| NEG_CNTRL_1day | 1.000000 | |
| NEG_CNTRL_5weeks | 0.034060 | * |
| 3_species_mix | 0 | ** |
| Grammatophora_marina_1week | 0 | ** |
| Grammatophora_marina_2weeks | 0 | ** |
| Grammatophora_marina_5weeks | 0 | ** |
| Achnanthes_brevipes_1day | 0 | ** |
| Grammatophora_marina_1day | 0 | ** |
| 10_species_mix | 0 | ** |
| 6_species_mix_settle | 0 | ** |
| 6_species_mix_settle_grow | 0 | ** |

Corresponding Kruskal-Wallis Test with Dunn's posthoc testing values:

All biofilm types tested, size of worms at 11 dpf (p-value cut-off = 0.05)

```
KruskalResult_Biofilm Age(statistic=99.51069604086854, pvalue=6.7015650
55929603e-20)
KruskalResult_Biofilm_Mix(statistic=177.57655619192968, pvalue=1.109695
7481070438e-35)
```

| | nbc_1d | nbc_5w | A.bre_1d | G.mar_1d | G.mar_1w |
|---|---|---|---|---|---|
| nbc_1d | 1.000000e+00 | 1.000000e+00 | 4.959869e-17 | 5.003905e-04 | 0.002048 |
| nbc_5w | 1.000000e+00 | 1.000000e+00 | 1.583681e-12 | 3.430884e-03 | 0.005270 |
| A.bre_1d | 4.959869e-17 | 1.583681e-12 | 1.000000e+00 | 5.709386e-04 | 0.106714 |
| G.mar_1d | 5.003905e-04 | 3.430884e-03 | 5.709386e-04 | 1.000000e+00 | 1.000000 |
| G.mar_1w | 2.047592e-03 | 5.270444e-03 | 1.067137e-01 | 1.000000e+00 | 1.000000 |
| G.mar_2w | 1.932183e-01 | 2.350559e-01 | 9.209193e-04 | 1.000000e+00 | 1.000000 |
| G.mar_5w | 8.096538e-04 | 2.458512e-03 | 2.115844e-01 | 1.000000e+00 | 1.000000 |
| Mix_3sp. | 7.197874e-27 | 1.396389e-19 | 9.603567e-01 | 9.824969e-10 | 0.000029 |
| Mix_6spS | 1.762342e-03 | 8.775185e-03 | 1.515784e-04 | 1.000000e+00 | 1.000000 |
| Mix_6spG | 6.211646e-22 | 5.715406e-16 | 1.000000e+00 | 1.234153e-06 | 0.002640 |
| Mix_10sp. | 2.157619e-13 | 5.921139e-10 | 1.000000e+00 | 3.446743e-02 | 1.000000 |

| | G.mar_2w | G.mar_5w | Mix_3sp. | Mix_6spS | Mix_6spG |
|---|---|---|---|---|---|
| nbc_1d | 1.932183e-01 | 0.000810 | 7.197874e-27 | 1.762342e-03 | 6.211646e-22 |
| nbc_5w | 2.350559e-01 | 0.002459 | 1.396389e-19 | 8.775185e-03 | 5.715406e-16 |
| A.bre_1d | 9.209193e-04 | 0.211584 | 9.603567e-01 | 1.515784e-04 | 1.000000e+00 |
| G.mar_1d | 1.000000e+00 | 1.000000 | 9.824969e-10 | 1.000000e+00 | 1.234153e-06 |
| G.mar_1w | 1.000000e+00 | 1.000000 | 2.852534e-05 | 1.000000e+00 | 2.640405e-03 |
| G.mar_2w | 1.000000e+00 | 1.000000 | 2.862498e-08 | 1.000000e+00 | 7.491399e-06 |
| G.mar_5w | 1.000000e+00 | 1.000000 | 8.197300e-05 | 1.000000e+00 | 6.330457e-03 |
| Mix_3sp. | 2.862498e-08 | 0.000082 | 1.000000e+00 | 1.432018e-10 | 1.000000e+00 |
| Mix_6spS | 1.000000e+00 | 1.000000 | 1.432018e-10 | 1.000000e+00 | 2.379755e-07 |
| Mix_6spG | 7.491399e-06 | 0.006330 | 1.000000e+00 | 2.379755e-07 | 1.000000e+00 |
| Mix_10sp. | 2.591310e-02 | 1.000000 | 4.414093e-02 | 1.190832e-02 | 1.000000e+00 |

| | Mix_10sp. |
|---|---|
| nbc_1d | 2.157619e-13 |
| nbc_5w | 5.921139e-10 |
| A.bre_1d | 1.000000e+00 |
| G.mar_1d | 3.446743e-02 |

```
G.mar_1w 1.000000e+00
G.mar_2w 2.591310e-02
G.mar_5w 1.000000e+00
Mix_3sp. 4.414093e-02
Mix_6spS 1.190832e-02
Mix_6spG 1.000000e+00
Mix_10sp.1.000000e+00
```

**Supplementary Figure S8**. Example light micrograph images for single and mixed microalgal biofilms used in settlement assays (Fig. 3)

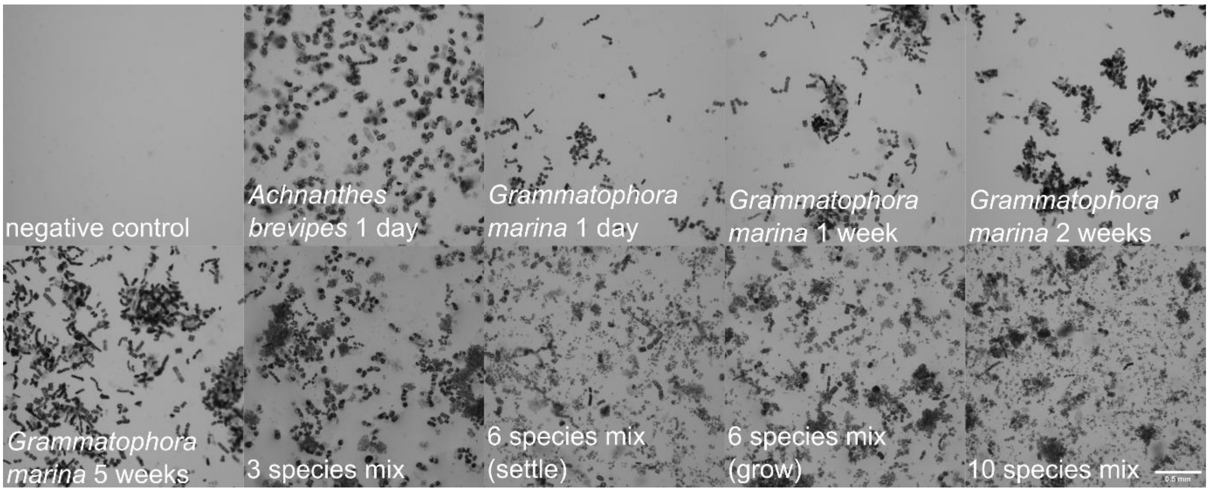


Fig. S8. Example light micrograph images of biofilms used in single and mixed microalgal species settlement assays. Biofilm composition name in bottom left corner. Scale bar 0.5mm.

**Supplementary Figure S9.** Example light micrograph images for postlarval growth at 11 days in response to different single and mixed species biofilms used in settlement assays (Fig. 3C)

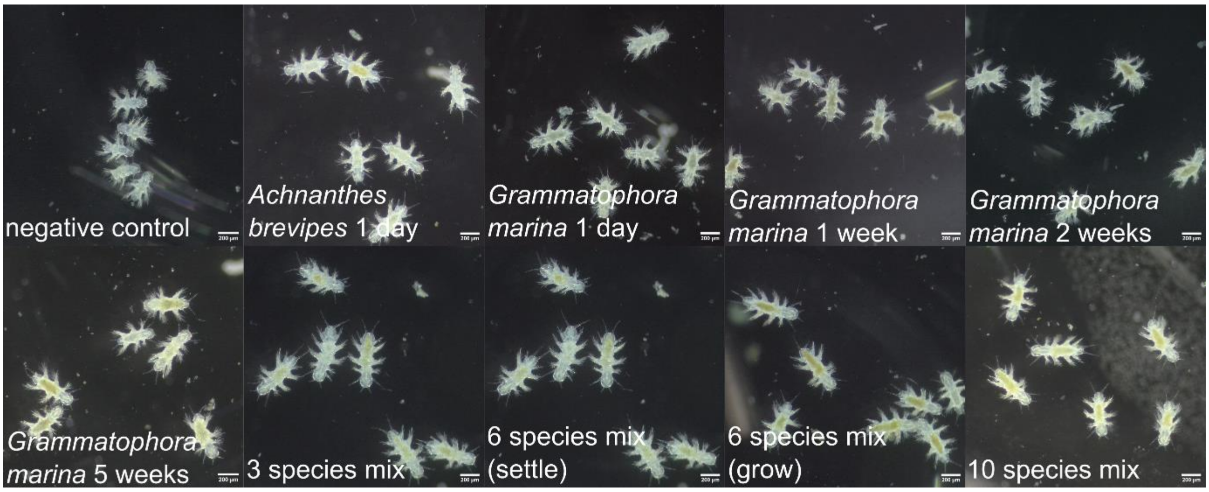


Fig. S9. Example light micrograph images of 11 days old postlarvae used in single and mixed microalgal species settlement assays. Biofilm composition name in bottom left corner. Scale bar 500 µm.

**Supplementary Table S11.** P-values of Mann-Whitney U Test on larval settlement in response to differently-treated *G.marina* microalgae biofilms (Fig. 4), at 24h and 48h post induction.

| Biofilm | 24h p-value | 48h p-value | Significance Code (*) |
|---|---|---|---|
| nbc | 1.000000 | 1.000000 | |
| G.mar_fil | 0.288487 | 0.935622 | |
| G.mar_EtOH | 0.024470 | 0.259818 | * |
| G.mar_boil | 0.008016 | 0.065081 | * |
| G.mar_50C | 0.030348 | 0.012592 | ** |
| G.mar_dry | 0.260658 | 0.008016 | * |
| G.mar | 0.004998 | 0.004998 | **** |
| G.mar_ab1 | 0.004998 | 0.004998 | **** |
| G.mar_ab2 | 0.004998 | 0.004847 | **** |

**Supplementary Figure S10.** Larval death during settlement assays in response to differently treated *G. marina* biofilms (Fig. 4A, B), at 24h and 48h post induction.

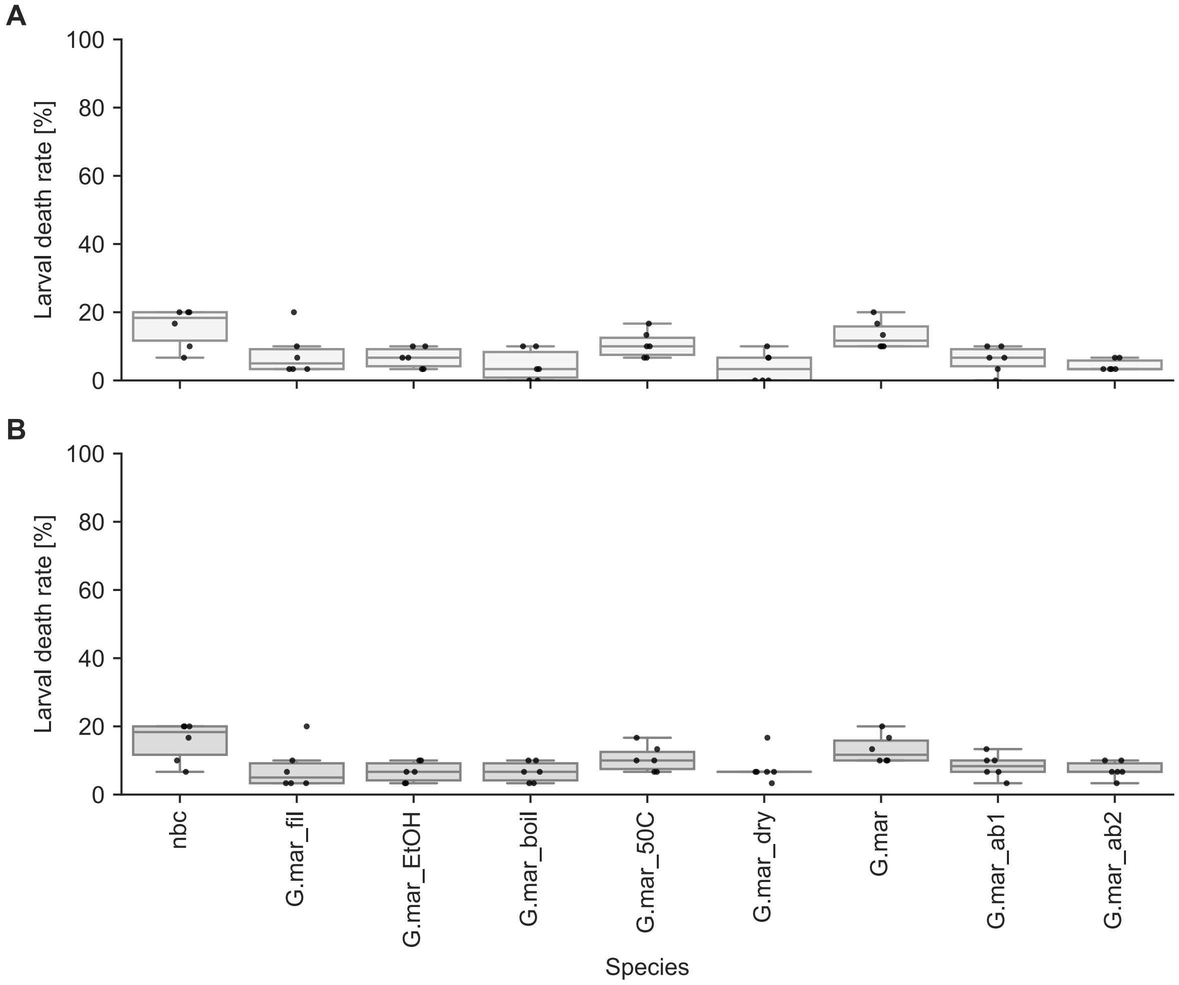


Fig. S10. Rates of death during settlement assays. Box plots with scatter plot overlay of (A) % larval death in response to differently treated *G. marina* biofilms after 24h and (B) 48h exposure. G.mar_fil = *G. marina* filtrate, G.mar_EtOH = EtOH-treated *G. marina*, G.mar_boil = boiled *G. marina*, G.mar_dry = *G. marina* 50℃ overnight (dry), G.mar_50C = *G. marina* 50℃ overnight (submerged), G.mar = 1 day old untreated *G. marina* biofilm, G.mar_ab1 = *G. marina* treated with antibiotics after biofilm formation, G.mar_ab2 = *G. marina* treated with antibiotics before and after biofilm formation, nbc = no biofilm control.

**Supplementary Table S12.** P-values of Mann-Whitney U Test on larval death in response to differently-treated *G. marina* biofilms, compared to a no biofilm control (nbc) (Fig. 3), at 24h and 48h post induction.

| Biofilm | 24h p-value | 48h p-value | Significance Code (*) |
|---|---|---|---|
| nbc | 1.000000 | 1.000000 | |
| G.mar_fil | 0.069855 | 0.069855 | |
| G.mar_EtOH | 0.026925 | 0.026925 | ** |
| G.mar_boil | 0.026925 | 0.018101 | ** |
| G.mar_50C | 0.140194 | 0.010194 | * |
| G.mar_dry | 0.029325 | 0.018101 | ** |
| G.mar | 0.505790 | 0.505790 | |
| G.mar_ab1 | 0.059922 | 0.027205 | ** |
| G.mar_ab2 | 0.031521 | 0.006345 | ** |

**Supplementary Figure S11.** Biofilm % coverage for the differently treated *G. marina* biofilms used in settlement assays (Fig. 4A, B)

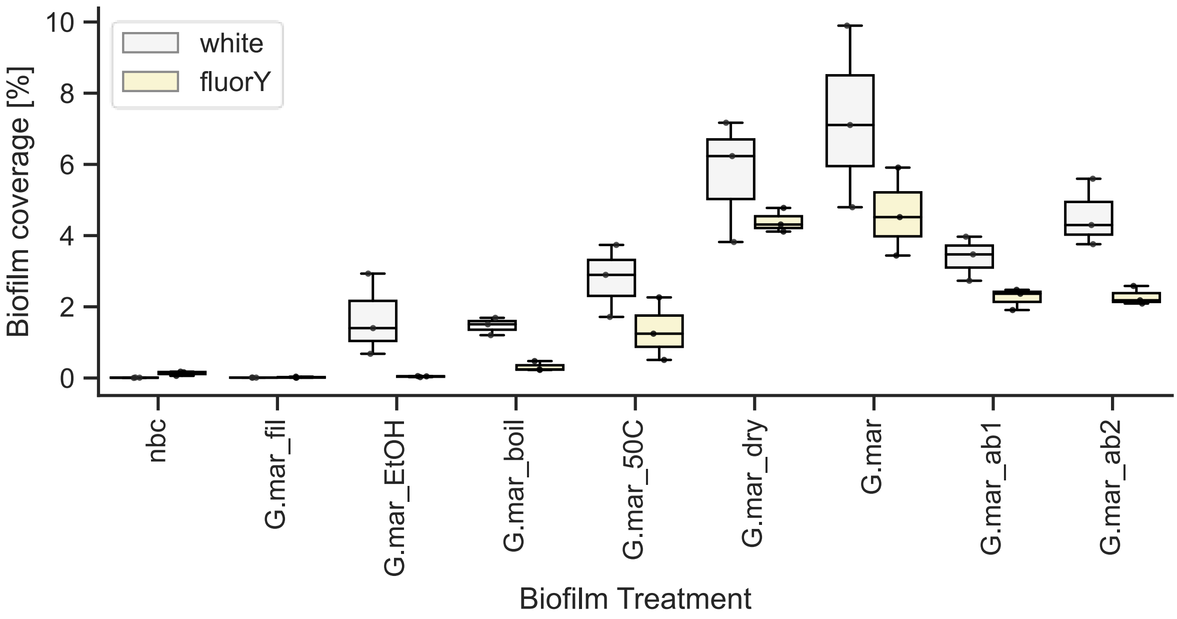


Fig. S11. Percent biofilm coverage during settlement on single or mixed microalgae species biofilms. Box plots of biofilm coverage [%] for differently treated *G. marina* biofilms at start of assay. G.mar_fil = *G. marina* filtrate, G.mar_EtOH = EtOH-treated *G. marina*, G.mar_boil = boiled *G. marina*, G.mar_dry = *G. marina* 50℃ overnight (dry), G.mar_50C = *G. marina* 50℃ overnight (submerged), G.mar = 1 day old untreated *G. marina* biofilm, G.mar_ab1 = *G. marina* treated with antibiotics after biofilm formation, G.mar_ab2 = *G. marina* treated with antibiotics before and after biofilm formation, nbc = no biofilm control. Coverage under white light represents total coverage including live and dead cells, coverage under fluorescent light with YFP filter represents live (viable) cell coverage (see also Fig. S12 for example images).

**Kendall's Tau correlation test results**, for correlation between biofilm density and larval settlement rate:

(1) All 8 biofilms tested, settlement at 24 hours post induction (hpi) under white light (all cells, live and dead)
Significance Result: statistic = 0.643, p-value = 0.031
(2) All 8 biofilms tested, settlement at 48 hours post induction (hpi) under white light (all cell, live and dead)
Significance Result: statistic = 0.286, p-value = 0.399
(3) All 8 biofilms tested, settlement at 24 hours post induction (hpi) under yellow fluorescence (viable/live cells)
Significance Result: statistic = 0.714, p-value = 0.014
(4) All 8 biofilms tested, settlement at 48 hours post induction (hpi) under yellow fluorescence (viable/live cells)
Significance Result: statistic = 0.357, p-value = 0.275

**Supplementary Figure S12**. Example light and fluorescent micrograph images for treated and untreated *Grammatophora marina* biofilms used in settlement assays (Fig. 4A,B)

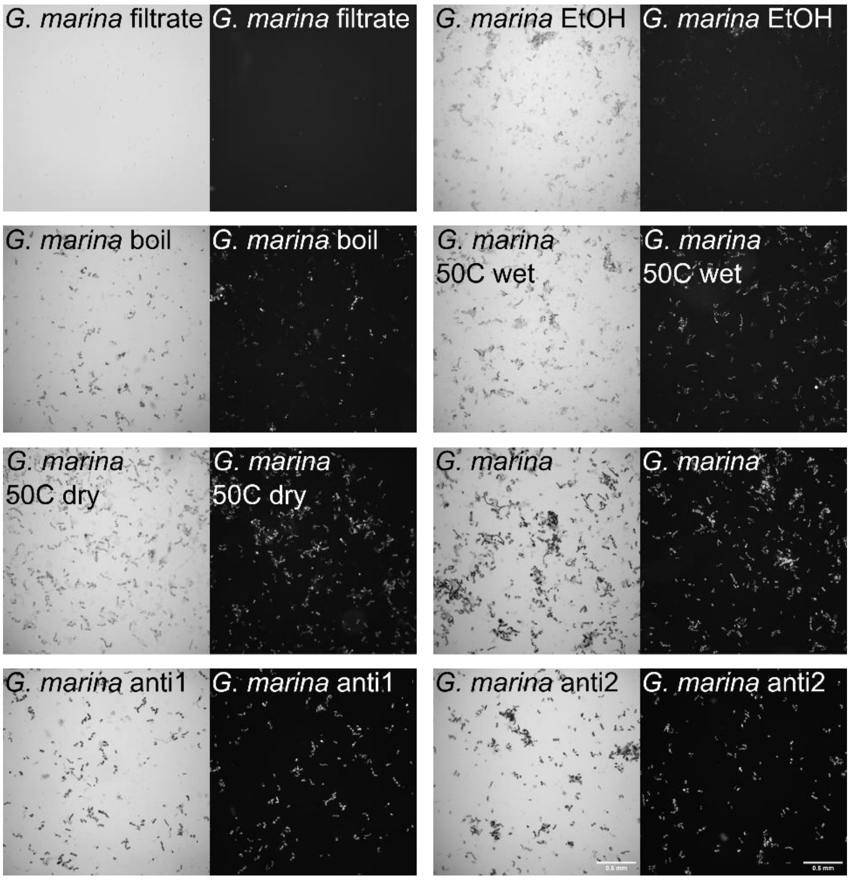


Fig. S12. Example light (left panel) and fluorescent (right panel) micrograph images of biofilms used in *G. marina* treatment settlement assays. Biofilm type in top left corner. Scale bar 0.5mm.

**Supplementary Table S13.** P-values of Mann-Whitney U Test on larval settlement of larvae of varying ages in response to 1 day old *G.marina* biofilm (Fig. 4C), at 4h, 24h and 48h post induction, compared to 8 days old larvae on a no biofilm control (nbc).

| Age of larvae | 4h p-value | 24h p-value | 48h p-value | Significance Code (*) |
|---|---|---|---|---|
| 8 days nbc | 1.000000 | 1.000000 | 1.000000 | |
| 2 days | 0.687363 | 0.285844 | 0.934568 | |
| 3 days | 0.226491 | 0.004772 | 0.004922 | **** |
| 3.5 days | 0.366480 | 0.004624 | 0.004922 | **** |
| 4 days | 0.020022 | 0.004847 | 0.004847 | ***** |
| 5 days | 0.008016 | 0.004922 | 0.004847 | ***** |
| 6 days | 0.004998 | 0.004772 | 0.004922 | ****** |
| 7 days | 0.004922 | 0.004847 | 0.004922 | ****** |
| 8 days | 0.004847 | 0.004772 | 0.004922 | ****** |

Note: % coverage of *G. marina* biofilms in larval age experiments = range 3.5 - 9.2%, nbc range = 0.002-0.008%. Median coverage *G. marina* biofilms in larval age experiments = 5.4%, nbc = 0.006%.

**Supplementary Figure S13.** Larval death during settlement assays of larvae of varying ages in response to 1 day old *G. marina* biofilm (Fig. 4C), at 4h, 24h and 48h post induction.

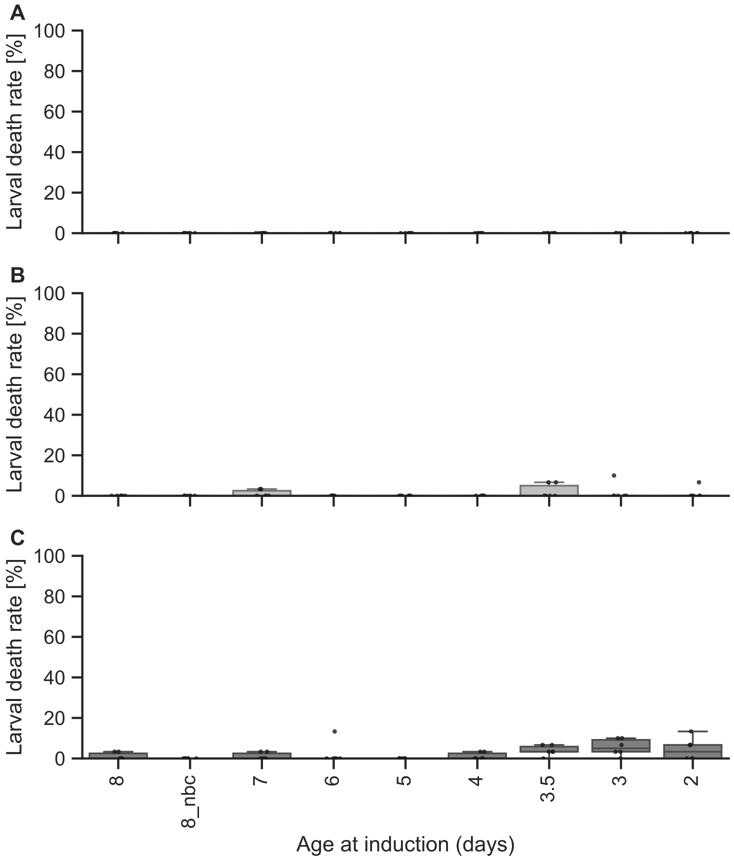

Fig. S13. Rates of death during settlement assays. Box plots with scatter plot overlay of % larval death in response to differently treated *G. marina* biofilms after (A) 4h, (B) 24h, and (C) 48h exposure.

**Supplementary Table S14.** P-values of Mann-Whitney U Test on larval death of larvae at different ages in response to 1 day old *G. marina* biofilms (Fig. 4C), at 4h, 24h and 48h post induction.

| Age of larvae | 4h p-value | 24h p-value | 48h p-value | Significance Code (*) |
|---|---|---|---|---|
| 8 days nbc | 1.000000 | 1.000000 | 1.000000 | |
| 2 days | 1.000000 | 0.404657 | 0.073140 | |
| 3 days | 1.000000 | 0.404657 | 0.009310 | * |
| 3.5 days | 1.000000 | 0.173945 | 0.008851 | * |
| 4 days | 1.000000 | 1.000000 | 0.173945 | |
| 5 days | 1.000000 | 1.000000 | 1.000000 | |
| 6 days | 1.000000 | 1.000000 | 0.404657 | |
| 7 days | 1.000000 | 0.173945 | 0.173945 | |
| 8 days | 1.000000 | 1.000000 | 0.173945 | |

**Supplementary Table S15.** P-values of Kruskal-Wallis Test with Dunn's posthoc testing on *Platynereis* total segment number at 30 days in response to different single species and mixed microalgal diets (Fig. 5).

| Diet | G.micro | Diatom | Mix_3sp. | Mix_5sp. |
|---|---|---|---|---|
| G.micro | 1.000000e+00 | 1.761871e-04 | 1.062757e-31 | 7.727772e-33 |
| Diatom | 1.761871e-04 | 1.000000e+00 | 6.957853e-10 | 7.847709e-12 |
| Mix_3sp. | 1.062757e-31 | 6.957853e-10 | 1.000000e+00 | 1.000000e+00 |
| Mix_5sp. | 7.727772e-33 | 7.847709e-12 | 1.000000e+00 | 1.000000e+00 |

G. micro = 1 species green microalgae, either *Tetraselmis suecica* or *Nannochloropsis salina*

Diatom = 1 species diatom, either *Phaeodactylum tricornutum*, or *Skeletoneman dohrnii*

Mix_3sp. = Mix of *Tetraselmis suecica*, *Nannochloropsis salina*, and *Phaeodactylum tricornutum*

Mix_5sp. = Mix of *Tetraselmis suecica*, *Nannochloropsis salina*, *Isochrysis galbana*, *Phaeodactylum tricornutum* and *Skeletonema dohrnii*.
